# Emergence of Slc11 clade MCb_gut_: a parsimonious hypothesis for the dawn of Lactobacillales in the gut of early vertebrates

**DOI:** 10.1101/2024.06.04.597488

**Authors:** M. FM Cellier

**Affiliations:** Centre Armand-Frappier Santé Biotechnologie, Institut National de la Recherche Scientifique (INRS), Laval, QC, H7V 1B7, Canada

**Keywords:** Lactic acid bacteria, proton-dependent Mn import, Slc11 carriers, AF2 modeling, conformational plasticity, in silico mutagenesis, phylogeny, evolution, vertebrates, stomach

## Abstract

The Lactobacillales (LB) stand apart among bacterial orders, using manganese (Mn) instead of iron to support their growth and swiftly ferment complex foods while acidifying their environment. The present work investigates whether a shift in the use of Mn could mark the origin of LB. Transmembrane carriers of the ubiquitous Slc11 family play key roles in LB physiology by catalyzing proton-dependent Mn import. In prior studies, the Slc11 clade found in LB (MntH Cb, MCb) showed both remarkable structural plasticity and highly efficient Mn uptake, and another Slc11 clade, MCg1, demonstrated divergent evolution coinciding with emergence of bacterial genera (e.g., *Bordetella*, *Achromobacter*). Herein, Slc11 clade MCb is subdivided in sister groups: MCb_ie_ and MCb_gut_. MCb_ie_ derives directly from Slc11 clade MCa, pointing an intermediate stage in the evolution of MCb_gut_. MCb_ie_ predominates in marine Bacillaceae, is more conserved than MCb_gut_, lacks the structural plasticity that typify MCb_gut_ carriers, and responds differently to identical mutagenesis. Exchanging MCb_ie_/MCb_gut_ amino acid residues at sites that distinguish these clades showed conformation-dependent effects with both MCb_ie_ and MCb_gut_ templates and the 3D location of the targeted sites in the carrier structure together suggest the mechanism to open the inner gate, and release Mn into the cytoplasm, differs between MCb_ie_ and MCb_gut_. Building on the established phylogeny for *Enterococcus* revealed that a pair of genes encoding MCb_gut_ was present in the common ancestor of LB, as MCb_gu1_ and MCb_gu2_ templates exhibit distinct structural dynamics properties. These data are discussed examining whether MCb ^+^ LB could emerge in the upper gut of early vertebrates (ca. 540 mya), through genome contraction and evolution toward Mn-centrism, as they specialized as gastric aids favoring stomach establishment in jawed vertebrates through bi-directional communication with host nervous, endocrine and immune systems.

## 1- Introduction

The Solute carrier 11 (Slc11) family of proton (H^+^)-dependent importers of Mn^2+^ and/or Fe^2+^ belongs to the APC superfamily and adopts the tridimensional (3D) LeuT fold [1], [2], [3]. This fold consists in tandem repeats of 5 transmembrane helices that display inverted transmembrane topology. The repeats fold intertwined and coexist in alternate 3D configurations that are exchanged as the carrier molecule switch between two main states, either outward open (OO) or inward open (IO). The absence of sequence similarity between the two halves of Slc11 carrier hydrophobic core implies asymmetric evolution which could favor directional substrate transport, i.e., H^+^-dependent import [4], [5].

Molecular evolutionary genetic and structural analyses revealed that Slc11 synapomorphy functionally distinguishes this protein family from the rest of the APC superfamily[6]. Slc11 synapomorphy comprises 11 (quasi) invariant amino acid (aa) residues forming a 3D network that articulates Slc11 substrate-selective carrier conformational exchange between the OO state and the IO state. Functional maintenance of this 3D network amid diverging Slc11 clades (e.g., bacterial MCb and MCg1) testifies of the resilience and evolvability of the molecular mechanism embedded through this synapomorphy in Slc11 architecture.

Besides, Slc11 mechanism of carrier coupling to the transmembrane protonmotive force (pmf), to catalyze cytoplasmic import of divalent metals such Mn and Fe (Me^2+^), demonstrated stepwise evolution in the form of serial synapomorphies that distinguish groups of Slc11 homologs found either in prokaryotes (the proton-dependent Mn transporters, MntH) or in eukaryotes (the Natural resistance-associated macrophage proteins, Nramp, Table S1).

MntH synapomorphies demonstrate successive sets of coevolutionary rate-shifts that progressively edified an extended ‘H^+^-network’ interacting with the pmf, from anaerobic bacteria (MntH B, before aerobiosis) to aerobic bacteria (MntH A) and to hyperthermophile TACK and Asgard Archaea (MntH H). This latter evolutionary intermediate likely shared a common ancestor with the first eukaryotic representative of the Slc11 family (Nramp precursor, Figure 1A, [7].

**Figure 1.**
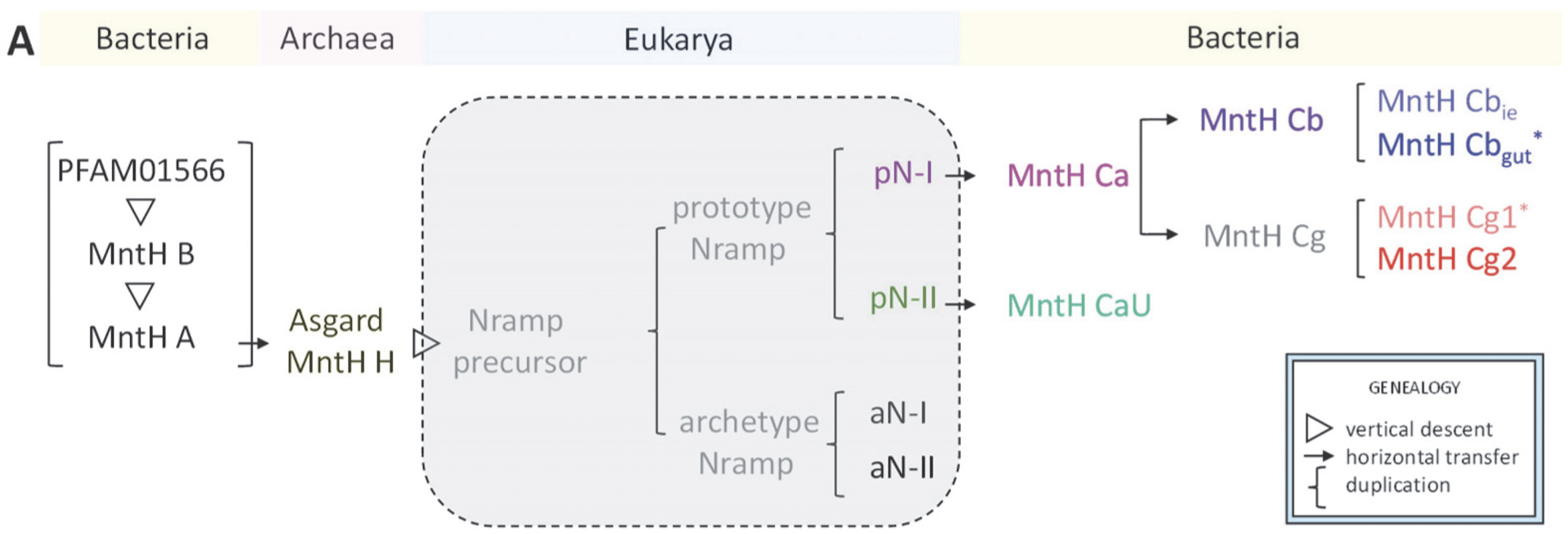
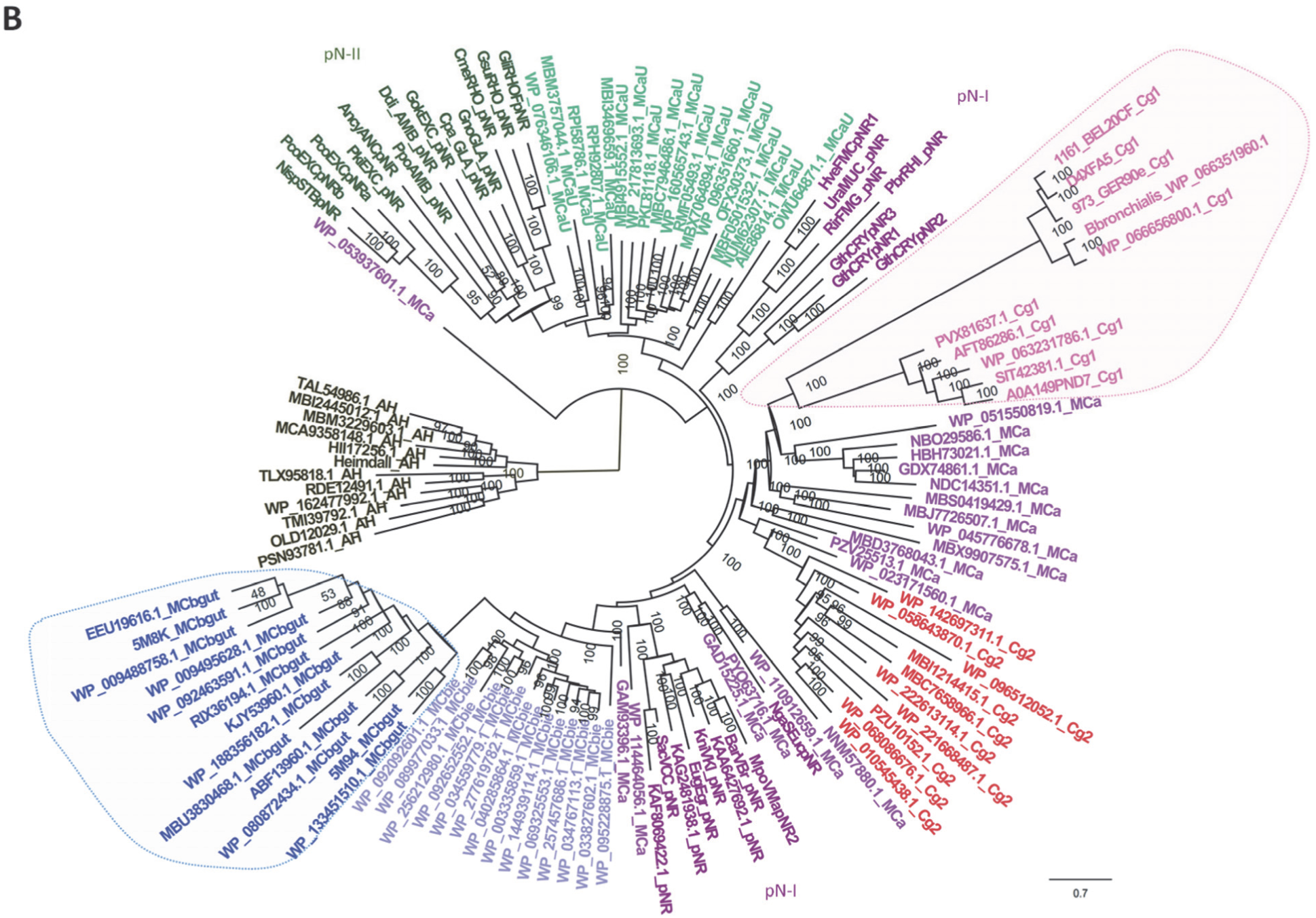
Phylogeny of bacterial Slc11 carriers of eukaryotic origin (MntH Ca (MCa), MCb, MCg and MCaU). **A**. Recapitulation of the current knowledge on evolution of the Slc11 family and proposition that MCb clade derived from MCa and comprises sister clusters, MCb_ie_ and MCb_gut_. Asterisks identify MC clades previously characterized as functionally divergent (highlighted in B). **B.** Identification of MCb_ie_ as possible evolutionary intermediate between MCa and MCb_gut_ clades. The tree presented was rooted using MntH H seqs from TACK and Asgard archaea. Phylogenetic reconstruction used 395 PI sites, the substitution model EHO, ML estimate of a.a. state frequency, free rate model of variation among sites with 15 categories. The statistical significance of each node is indicated together with the scale (nb substitution per site). This phylogeny demonstrates partition of eukaryotic pN-I and pN-II parologs, and the common ancestries of pN-II and MCaU on one hand, and pN-I and MCa, MCb, MCg on the other hand. MCb and MCg clades apparently derived from distinct MCas. The topology of MCb group suggests divergent evolutionary trajectories for MCb_ie_ and MCb_gut_, as previously observed in MCg1 subgroup [6].

Slc11 H^+^-network further evolved in eukaryotes, during transition from first to last eukaryotic common ancestor (FECA to LECA), yielding two types of Nramp: prototype Nramp, mainly found in primitive eukaryotes, and archetype Nramp, predominant in multicellular organisms of both animal and vegetal realms. Coupled to phagocytosis, archetype Nramp can restrict bacterial survival by depleting the acidified phagosomal milieu of Me^2+^, such as Mn and Fe [8]. Arguably, evolution of Slc11 mechanism of carrier coupling to the pmf resulted in functional antagonism, yielding eukaryotic Nramp carriers that import Me^2+^ more efficiently than their bacterial counterparts (MntH).

However, both types of prototype *Nramp* genes (*pN-I* and *pN-II*, Table S1) were independently transferred horizontally towards bacteria. Therefore, bacterial MntH are polyphyletic [9], either of prokaryotic origin (MntH B, MntH A, MntH H) or of eukaryotic ancestry (MntH C). Alphafold (AF) [10] / Colabfold (CF) [11] 3D modeling of native MntH carriers of eukaryotic origin, from the sister clades MntH Cb (MCb) and MntH Cg1 (MCg1) Figure 1), coupled to modeling of phylogeny-informed site-directed mutants, demonstrated paths of divergent evolution which affected both the carrier shape and functional plasticity [6].

A hallmark of MCb carriers is their conformational flexibility, and protein sequences from distinct clades (>80% sequence identity) yielded 3D models representing discrete conformers. Regarding MCg, a mutation altering both the H^+^-network in transmembrane helix 4 (h4) and carrier shape contributed to found MCg1 clade, and further divergent epistasis within this clade yielded an apparently functional neovariant, whose emergence coincided with novel bacterial genera (*Bordetella* and *Achromobacter*). This unexpected result suggested that, by analogy, emergence of MCb carrier might correspond as well to the rise of other bacterial genera.

Indeed, MCb is the only Slc11 type found in the order Lactobacillales (LB), and the majority of known MCbs are encoded by LB genomes. In contrast, the close relatives Bacillales may harbor either *mntH A*, *mntH Ca* or *mntH Cb* genes [7]. The extraordinary flexibility and functional plasticity of MntH Cb 3D models may relate to high carrier efficiency, able to support LB’s *mntH*-dependent competitive exclusion of other organisms [12], [13].

It is perhaps no coincidence that LB stand out as a bacterial order that relinquished heme-based metabolism to become Mn-centric microorganisms. As Mn excess can jeopardize iron-based metabolic functions, high efficiency Mn import seems particularly suited to cells notably lacking iron-cofactored enzymes for respiration, antioxidant defenses and the tricarboxylic acid cycle [14], [15], and which rely on Mn instead of Fe for growth [16], [12], [17]. Based on available data it can therefore be hypothesized that emergence of MCb clade was concomitant to the dawn of LB.

The consensus on LB origin indicates they derived from a Bacillale ancestor through ∼1/3 genome reduction and by adapting to some microaerated/anaerobic nutrient-rich environmental niche [14], [15]. Given that LB are both notoriously proficient at predigesting (fermenting) complex food and common residents of animal gut [18], [19], it seems plausible that LB could originate there. LB are demonstrated residents of the gut of various vertebrates (fish, reptiles, birds, mammals) and a landmark study proposing the genus *Enteroccoccus* arose more than 425 million years ago (mya), as animals colonized land [20], suggests LB common ancestor could evolve earlier, during the Ediacaran-Cambrian transition for instance.

To test the hypothesis that emergence of (Slc11) MCb clade may coincide with LB origin, a sequence of possible evolutionary events linking MCa and MCb clades were established, and their functional implications tested in silico using AF/CF 3D modelling. Several questions were addressed: Is there a group of MntH Cb seqs that display common properties identifying them as possible intermediates (hereby named MCb_ie_, for MCb environmental intermediate) between MntH Ca and the rest of MCb clade (hereafter named MCb_gut_)? Do MCb ^+^ microorganisms form a defined Bacillale taxon and/or share a common habitat? How do MCb_ie_ and MCb_gut_ in silico structural dynamics compare? Do they support a distinct mechanism of Mn import in LB? Does MCb_gut_ emergence predate the origin of Enterococci and what was the *mntH* Cb_gut_ gene complement of the common ancestor of LB?

The results presented define the Slc11 clades MCb_ie_ and MCb_gut_ which together form the phylogroup MCb. MCb_ie_ predominates in marine Bacillaceae and is closely related to both its ancestor clade, MCa, and the common precursor of MCb_gut_. Regarding MCb_gut_, it is in fact a pair of genes, encoding MCb_gu1_ and MCb_gu2_, that typifies LB. The 3 MCb clades MCb_ie_, MCb_gu1_ and MCb_gu2_ vary in their evolutionary conservation and conformational properties as opening of the carrier inner gate apparently evolved in the transition from MCb_ie_ to MCb_gut_. These data are discussed relating the emergence of MCb ^+^ LB and development of the gastro-intestinal tract of early vertebrates.

## 2- In Silico Results

### 2.1. *P. halotolerans* A0A1I3CNB9 identifies the clade MCb_ie_, intermediate between MCa and MCb_gut_

Prior work suggested that reevaluating the phylogenetic structure of MCb clade may delineate a putative precursor lineage (MCb_ie_) from LB’s MCbs (MCb_gut_): NCBI sequence wp_092092601.1 from the Carnobacteriaceae *Pisciglobus halotolerans* (Uniprot ID: A0A1I3CNB9), representing a cluster of sequences (seqs>70% aa id), demonstrated both MCb most basal branching node and relatively little specific divergence compared to the rest of MCb clade [7]. Detailing the phylogenetic relationships of A0A1I3CNB9 clade, i.e., MCb_ie_, amid eukaryotic prototype Nramps and other bacterial MntH C clades (Figure 1B) confirmed both its basal position relative to the rest of MCb homologs, i.e., MCb_gut_, and the relative sequence conservation of its members.

MCb_ie_ conservation contrasts with the diversity of MCa clade, which comprises both the closest known relatives of prototype Nramp-I (pN-I) and candidate precursors of both MCb and MCg clades [7]. The phylogenetic position of MCb_ie_ identifies an evolutionary intermediate between MCa and MCb_gut_ (Figure 1 and Figure S1). MCb_ie_ clade topology shows 2 subsets (Figure S1A): a ‘crown’ formed by conserved sequences from core marine Bacillales (Bacillaceae, Paenibacillaceae) plus more divergent subsets from other (Lacto)bacillales spp. that exhibit relatively reduced genomes.

MCb_ie_ is most prevalent in Bacillaceae (45 seqs, 95% id cutoff, Figure S1A) and MCb ^+^ genera represent about 10% of the Bacillaceae family (Figure S1B). MCb_ie_ phylogeny segregates several established genera of Bacillaceae, a pattern seemingly compatible with vertical inheritance of *mntH* Cb_ie_. Yet both Bacillaceae and Planococcaceae encode predominantly the more ancestral MntH A (MA) protein, which contrasts with Paenibacillaceae wherein MntH Ca (MCa) predominates (205 seqs, 95% id cutoff). Neither Listeriaceae [21], [22], [23] nor Carnobacteriaceae (LB) harbor any MA coding gene, and contrarily to Paenibacillaceae, MCb (MCb_ie_ or MCb_gut_) is enriched in both these families. In comparison, Lactobacillaceae possess exclusively MCb_gut_ (MCb_gu1_ and/or MCb_gu2_, detailed in section 2.6).

The topology of MCb_ie_ ‘crown’ group shows clusters of seqs from Paenibacillaceae that diverge from clusters of seqs from Bacillaceae (Figure S1A). Besides, only spp. of Paenibacillaceae (e.g., *Paenibacillus*: NZ_CP019717, NZ_CP020327 and *Brevibacillus*: CP048799, CP139435) were found harboring both genes encoding MCa and MCb_ie_, including some *Paenibacillus* isolates wherein MCb_ie_ appears inactivated (NZ_CP019794, NZ_CP020557). Together with the large genome size of MCb_ie_^+^ Paenibacillaceae spp., the data support that MCb_ie_ could originate in Bacillaceae and subsequently spread among Paenibacillaceae spp. Similarly, *mntH* Cb_ie_ gene transfers toward spp. showing genome erosion (e.g., Listeraceae and Carnobacteriaceae) may account for the relative divergence of their encoded MCb_ie_ proteins.

Phylogenomic analyses [24], [25] support this interpretation, showing marine MCb_ie_^+^ Bacillaceae, e.g., *Heyndrickxia* and *Lederbergia* genera, as more likely candidate precursors of LB than the MA^+^ *Bacillus* spp. (*B. subtilis* and *B. anthracis*, Figure 2 and Figure S2). Staphylococcaceae and Listeriaceae are known as highly derived Bacillales families of Fe-dependent bacteria [26], [27] and both lack MA encoding genes.

**Figure 2.**
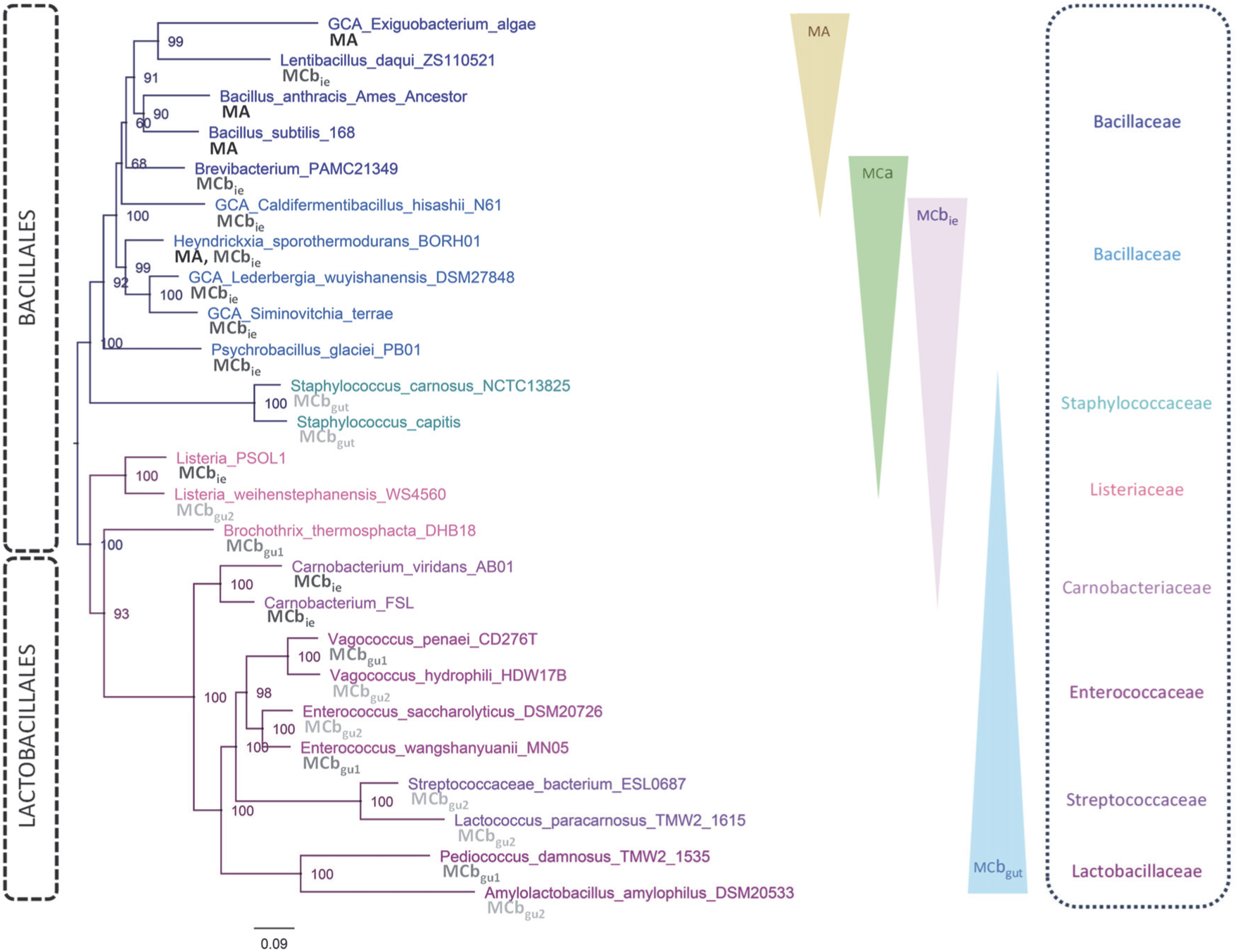
Bacillales and Lactobacillales phylogeny. Based on the VBCG pipeline [24] using select *mntH* encoding genomes plus IQ-Tree reanalysis (EX-EHO+R6) of the VBCG concatenated alignment. Species names are color-coded at family-level (indicated, *right*). *mntH*-encoded isotype is indicated per genome (*bold*). Appreciation of *mntH* gene flows according to the present hypothesis is schematized using colored arrowheads.

Altogether the data presented clarify the emergence of the Slc11 clade MCb from its precursor MCa (itself derived from eukaryotic pN-I, Figure 1A). An intermediate step in this process produced the sister clades MCb_ie_ and MCb_gut_, respectively predominant in marine Bacillaceae, such as *Heyndrickxia* and *Lederbergia*, which are aerobic, and in anaerobic LB. These results provide molecular insight into the hypothetical evolutionary transition relating MA^+^ Bacillales to MCb ^+^ LB through a marine MCb ^+^ Bacillaceae intermediate.

### 2.2. AF/CF 3D modeling of native MCb_ie_ and MCb_gut_ carriers differs significantly

Multiple alignment of diverse sequences from each three groups MCa, MCb_ie_ and MCb_gut_ demonstrate 29 membranous sites that coevolved in the transition between MCa and MCb, i.e., coevolved mutations fixed in the common ancestor of MCb_ie_ and MCb_gut_ (Figure 3AB). Also, sequence variations in extra membranous loops of the carrier molecule show several instances of selective conservation between MCa and MCb_ie_ vs MCb_gut_ (e.g., l 4/5, l6/7, l7/8, l8/9 and l9/10, Figure S3), indicating that the common precursor of MCb clade evolved directly from MCa.

**Figure 3.**
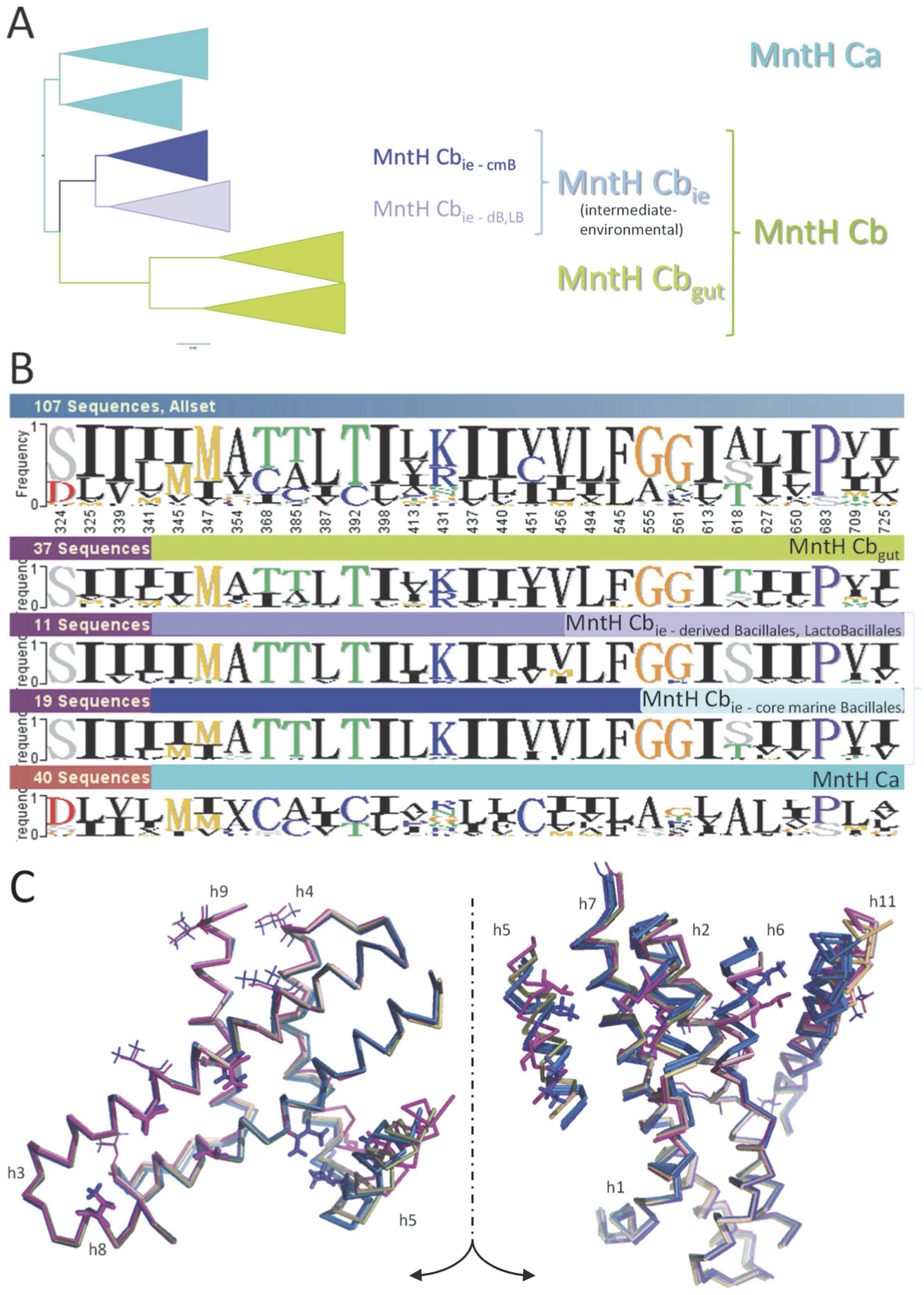
Structural evolution of MntH C in transition from MCa to MCb. **A**. Similarity clusters highlight relationships between the groups of sequences analyzed in B. **B.** Phylo-mLogo display of the site-specific evolutionary rate-shifts that discriminate aligned MCb seqs (MCb_ie_ plus MCb_gut_) from MCa seqs. **C.** 3D location of the coevolutionary rate-shits presented in B. Superposed models (Cα ribbon) of previously characterized MCb conformers (AF-Q5HQ64, magenta, CF-Q5HQ64, light pink, CF-Q5HQ64_VY-YN-228T, dark blue; CF-A0A380H8T1, light green, CF-A0A380H8T1_VN223T, light blue; CF-WP_002459413, light orange, CF-WP_002459413_VN224T, marine blue [6] allow visualizing site motion during carrier cycle. MCb architecture is presented after split of transmembrane helix 5 (h5) and opposite rotation of each half to display views from the center of the molecule. Transmembrane helices are numbered, and the extra membranous loops were omitted for clarity.

3D display of these 29 coevolved sites between MCa and MCb, on superposed MCb_gut_ models representing both the OO and IO states (cracked-open representation segregating half segments of helix 5 (h5), Figure 3C), shows accumulation along h5. The tilt of h5 modeled during MCb_gut_ carrier conformation switch was shown to mobilize separate communities of networked residues, involving one or the other half of h5, and each contributing to open the inner gate of the carrier toward the cell interior [6]. The 29 (MCa/MCb) rate-shifted sites segregate in distinct areas involving either h5C, h7C, h1b, h2N, h6a and h11C (right panel) or h5N, h4C and h8N plus additional sites scattered across the ‘hash module’ which hosts a large part of Slc11 H^+^-network (left panel).

These data indicate that evolution of the precursor of MCb clade directly from MCa involved a set of coevolved sites. These evolutionary coupled mutations, common to both MCb_ie_ and MCb_gut_, may form a clade-specific (MCb) synapomorphy. Their accumulation in local areas of the 3D structure that involve one half or the other of h5, whose tilting during MCb_gut_ carrier cycle connects outer gate locking and inner gate opening [6], suggests these processes evolved in the transition from MCa to MCb common ancestor.

The 3D modeling properties of MCb_gut_ templates differ from other MntH C clades, apparently due to intrinsic carrier plasticity and the presence of flexible loops [6, 7]. Based on the candidate MCb synapomorphy (Figure 3BC) one could expect that AF2/CF models of native MCb_ie_ carriers would resemble known MCb_gut_ structures either modeled or solved. However, 3D modeling of MCb_ie_ templates differed significantly (Figure 4).

**Figure 4.**
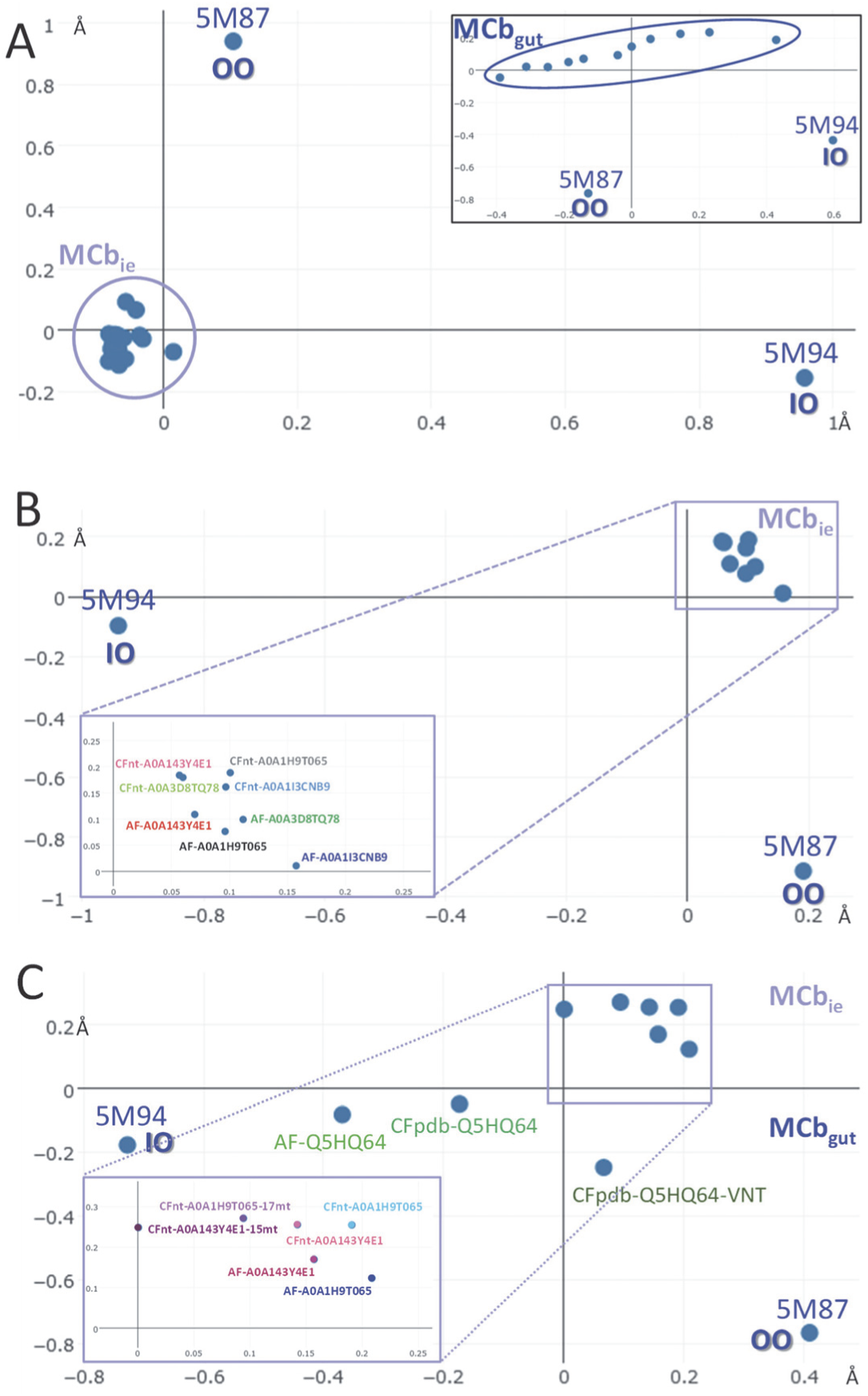
3D modeling (AF2 or CF) demonstrates structural difference between MCb_ie_ and MCb_gut_. **A**. Structural correspondence established by pairwise alignments of sampled AF2 models of MCb_ie_ carriers and the solved reference structure of MCb_gut_ carriers, either outward-open (5M87, [28]) or inward-open (5M94, [29]). *Inset*, array of native MCb_gut_ models Cellier [6], [7]. **B.** Structural correspondence between AF2 and CFnt models of 4 MCb_ie_ templates, detailed enlarged. **C.** Impact of compound mutagenesis targeting a set of coevolved sites that discriminate MCb_ie_ from MCb_gut_ (presented in Figure 5A) on CFnt 3D modeling of select MCb_ie_ templates. Additional reference models include MCb_gut_ Q5HQ64 AF2 and CFpdb models (IO) and the Q5HQ64 VNT mutant (OO, CFpdb, [6]).

3D correspondence of available Uniprot/AF2 MCb_ie_ models showed very similar predicted structures that did not clearly segregate with either the OO state or the IO state of solved structures of MCb_gut_ homologs (Figure 4A). This result contrasts with the array of functional conformers that AF2 predicted using MCb_gut_ sequences (Figure 4A, inset).

Examining models obtained for 4 representative MCb_ie_ templates (∼70% pairwise aa id) shows they do not represent clearly distinct OO or IO candidate conformers (Figure 4B), and their sequence relationships (cf Figure 1B, i.e., between WP_256212980.1 (NCBI proxy for Uniprot A0A143Y4E1), WP_092092601.1 (A0A1I3CNB9), WP_092652552.1 (A0A1H9T065) and WP_277619782.1 (proxy for A0A3D8TQ78)) suggests MCb_ie_ phylogeny does not influence AF2 or CFnt predictions. The 2 sequences that produced similar CFnt models, apparently closer to MCb_gut_ IO state, were selected for further analysis (A0A143Y4E1 and A0A1H9T065, both from Carnobacteriaceae spp., *Trichococcus palustris* and *Isobaculum melis*, respectively).

The topology of MCb clade (MCb_ie_ and MCb_gut_) resembles the previously characterized MCg1 clade ([6] and Figure 1B): both subdivide in sister groups that differ in relative extent of divergence. For MCg1 clade, this situation was correlated with i) founding mutations that introduced intrinsic structural constraints and ii) reversion to less constrained model structure as a result of extensive epistatic divergence.

Regarding MCb_gut_, divergence from MCb common ancestor produced remarkably flexible and intrinsically dynamic structures, whose predicted conformations reflected phylogenetic diversity (Figure 4A, inset). In contrast, the 3D models obtained using relatively diverse MCb_ie_ seem structurally constrained (Figure 4B), suggesting that MCb_gut_ could evolve out of epistatic divergence from (its common ancestor with) MCb_ie_.

### 2.3. Evolution of MCb carrier cycling between MCb_ie_ and MCb_gut_

To examine this possibility functional evolution between MCb_ie_ and MCb_gut_ was investigated by comparing AF/CF 3D modeling of native and mutant templates targeted at sites that underwent coevolutionary rate-shifts in transition from MCb_ie_ to MCb_gut_ (putative MCb_gut_ synapomorphy). 15 or 16 mutations that distinguish MCb_gut_ from MCb_ie_ were cointroduced into the sequences A0A143Y4E1 and A0A1H9T065, respectively (71.5% aa id, Figures 4C, 5A & Table S2). This approach aimed at testing the impact on MCb_ie_ predicted structure of 17 comutations that typify instead the MCb_gut_ clade.

As a result, CFnt modeling of the compound mutant A0A143Y4E1_15mt (Table S2) predicted a structure closer to the IO state of MCb_gut_ (Figure 4C). This suggested MCb_gut_ synapomorphy may signify functional determinants, as the corresponding residues map to 3D areas either partially overlapping with MCb synapomorphy (Figure 5A and Figure 3BC, respectively) or specific to MCb_gut_ (e.g., l 7/8). In addition, while native MCb_ie_ A0A143Y4E1 CFnt model cannot fully superimpose onto MCb_gut_ IO CFpdb model (Q5HQ64_WFLF, [6]) the predicted structure for the compound mutant A0A143Y4E1_15mt does so almost perfectly (Figure 5B).

**Figure 5.**
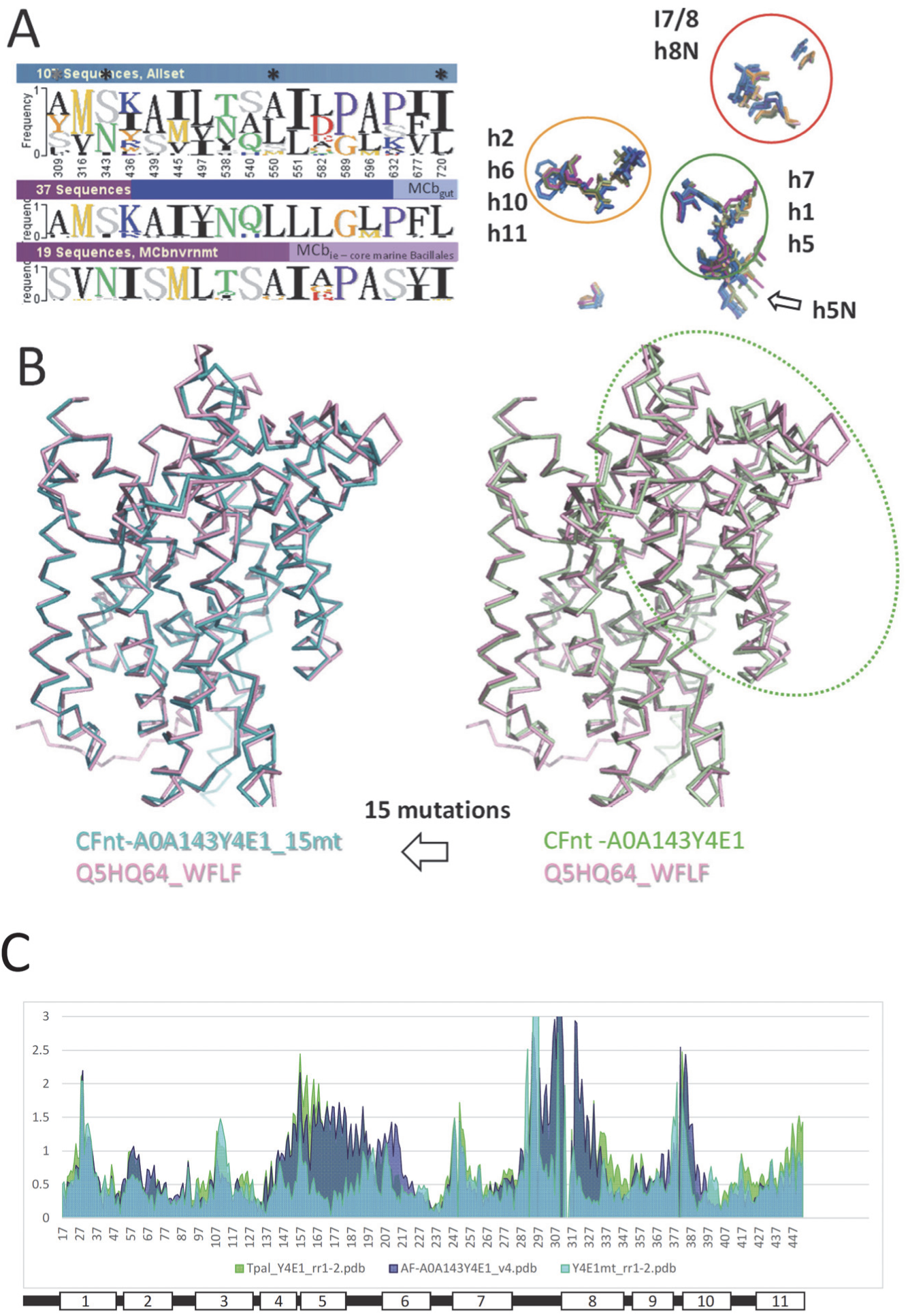
Structural reorganization of MCb_ie_ A0A143Y4E1 after exchanging 15 residues that distinguish MCb_ie_ and MCb_gut_ clades. **A**. 17 type ii evolutionary rate-shifts identified by comparing aligned representative seqs of MCb_ie_ (core marine Bacillales) and MCb_gut_ clades. Left panel, Phylo-mLogo display, right panel, 3D location on MCb_gut_ models. **B.** 3D superposition onto native MCb_gut_ CFpdb model (Q5HQ64_WFLF, inward open) of MCb_ie_ A0A143Y4E1 CFnt models predicted for the native (right) and compound mutant (A0A143Y4E1_15mt, left) templates. A dotted line shows local deviation between the structures compared. **C.** Per residue root mean square deviation (pRMSD) from MCb_gut_ inward open predicted structure (Q5HQ64_WFLF, CFpdb) of models obtained for MCb_ie_ A0A143Y4E1 templates either native (AF2, CFnt) or after compound mutagenesis (A0A143Y4E1_15mt, CFnt).

These data suggest that a select portion of MCb_gut_ carrier, which contributes to inner gate opening [6], was rearranged compared to MCb_ie_. Per residue root mean square deviation (pRMSD) from MCb_gut_ IO CFpdb model (Q5HQ64_WFLF) shows the segments rearranged in MCb_ie_ compound mutant include l1/2 & h2, h4-h6, l 7/8 & h8, l 9/10 & h10 (Figure 5C). These structural rearrangements extend beyond the areas targeted for mutagenesis (Figure 5A), implying the targeted sites may directly mediate MCb carrier conformation change.

Investigating this further, the reciprocal mutational approach was applied to MCb_gut_ homologs from distinct clades (>80% aa id) that produce discrete native conformers, either OO or IO, or in-between ([6] and Figure 6A), to determine how MCb_ie_-specifying residues affect MCb_gut_ predicted structure. This yielded two types of results: i) for templates whose predicted native conformers adopt the IO state, the 17 mutations introduced resulted in the prediction of OO conformers, a result similar to that obtained by targeting instead 5 sites of Slc11 synapomorphy ([6] and Figure 6AB); ii) considering templates whose predicted native conformers adopt the OO state or in-between state, identical compound mutagenesis had comparatively little or no effect.

**Figure 6.**
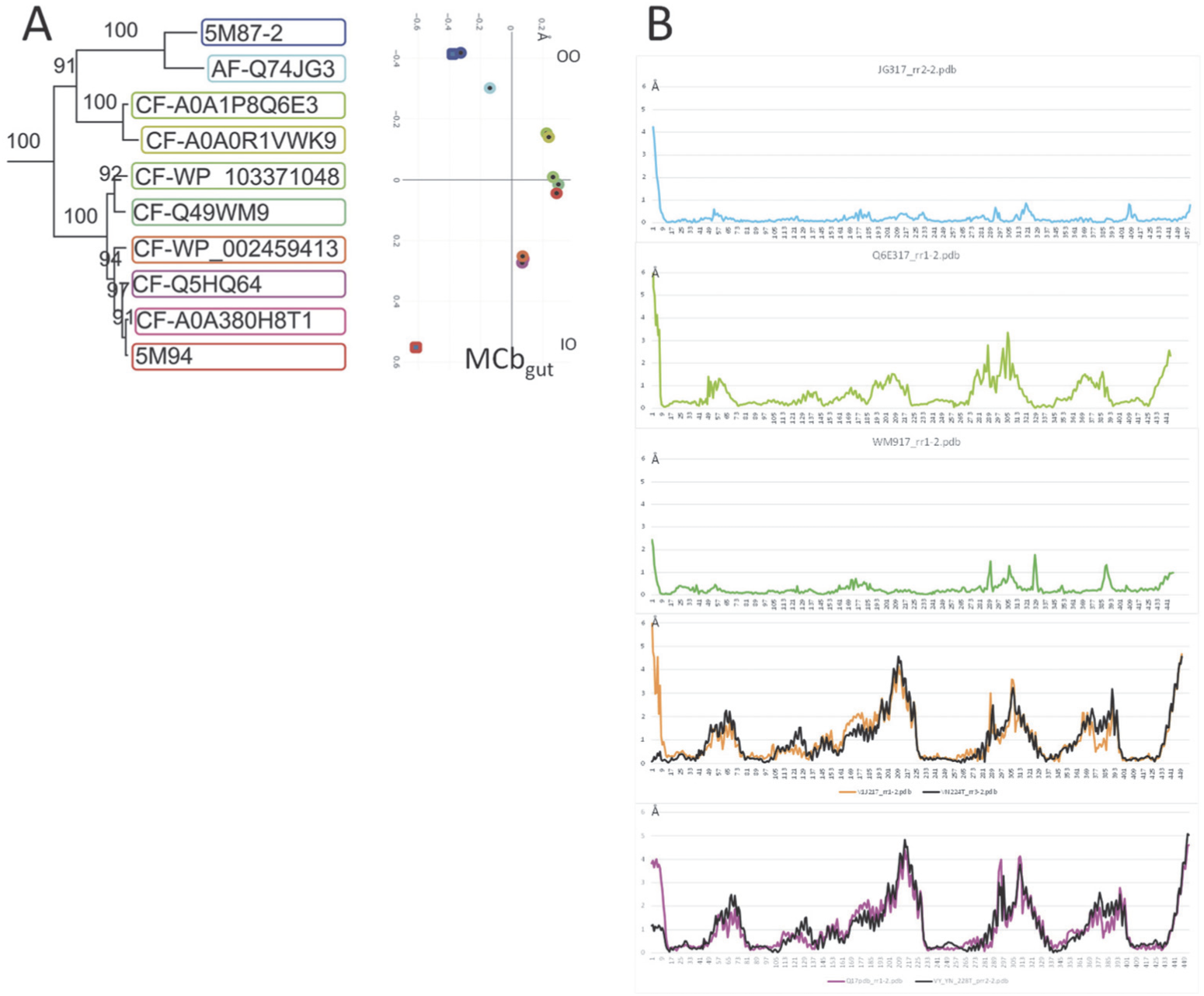
Conformation-dependent impact on MCb_gut_ CFpdb modeling of substituting 17 residues that discriminate MCb_ie_ and MCb_gut_ (reciprocal mutagenesis cf Figure 5). **A**. MCb_gut_ phylogeny was previously shown to influence the conformers modeled by CFpdb [6]. **B.** Co-introduction of 17 mutations in select MCb_gut_ templates shows conformation-dependent effects. Native CFpdb models were used as reference to determine the pRMSD of each respective mutant model. From top to bottom: Q74JG3, A0A1P8Q6E3, Q49WM9, WP_002459413, Q5HQ64. The pRMSD of WP_002459413 VNT and Q5HQ64 VNT mutant models [6] is also shown to visualize the expected deviation when comparing inward open and outward open conformers.

The conformational effects of the reciprocal mutageneses performed were both asymmetric and specific, favoring an MCb_gut_-like IO state for the MCb_ie_ A0A143Y4E1_15mt mutant while triggering MCb_gut_ conformation switch from IO state to OO state but showing marginal effects on alternate MCb_gut_ carrier conformational states (OO and in-between). Together these results demonstrate the functional impact of exchanging a set of coevolved residues that distinguish MCb_gut_ from MCb_ie_, showing conformation-dependent effect on MCb_gut_ carrier 3D modeling.

### 2.4. Mutagenesis of Slc11 synapomorphy supports functional divergence of MCb_gut_ and MCb_ie_

Comparative analysis of CF modeling sensitivity to mutagenesis targeting Slc11 synapomorphy has proven useful to decipher functional evolution between MCb and MCg1 clades and within MCg1 clade [6]. Introduction of the canonical VNT mutation (h6 VY, h3 YN, h6 T) into templates from both clades resulted in the conversion of native IO conformers into OO mutant conformers. However, the respective sensitivity to either h3 YN and/or h6 VY ^+^/_-_ h6 T mutations varied between clades.

Applying the VNT mutation to MCb_ie_ A0A143Y4E1 produced the expected CFnt modeling result which predicted an OO conformer like the reference MCb_gut_ Q5HQ64-VNT [6], indicated by comparable broad scale rearrangement (Figure 7A) and quasi-superimposable structures (Figure 7B). In contrast, none of native A0A143Y4E1 predicted structures (using CFnt or AF2) nor CFnt model for A0A143Y4E1_15mt fitted alike MCb_gut_ OO model conformer. Hence, mutating 5 sites of Slc11 synapomorphy induced a conformational response of MCb_ie_ A0A143Y4E1 like those observed before with MCb_gut_ and MCg1 clades.

**Figure 7.**
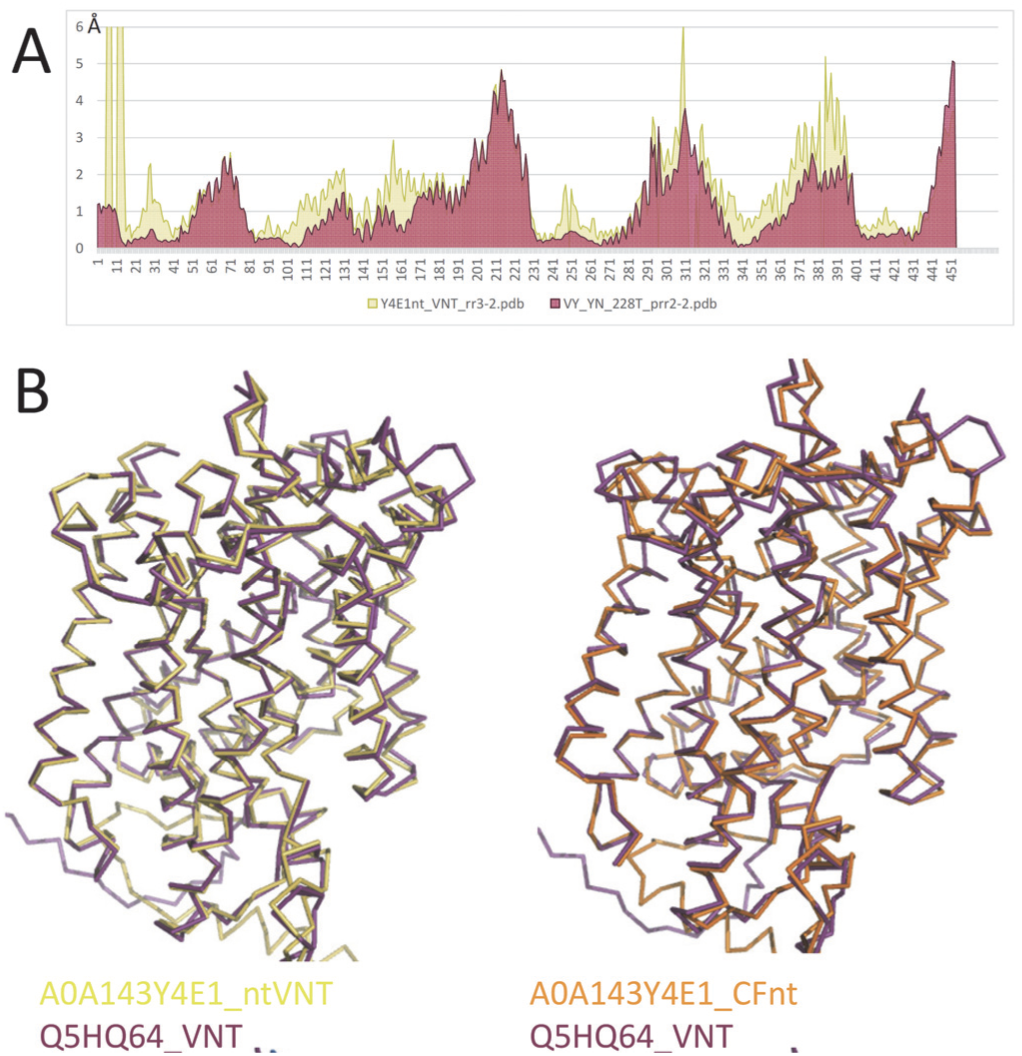
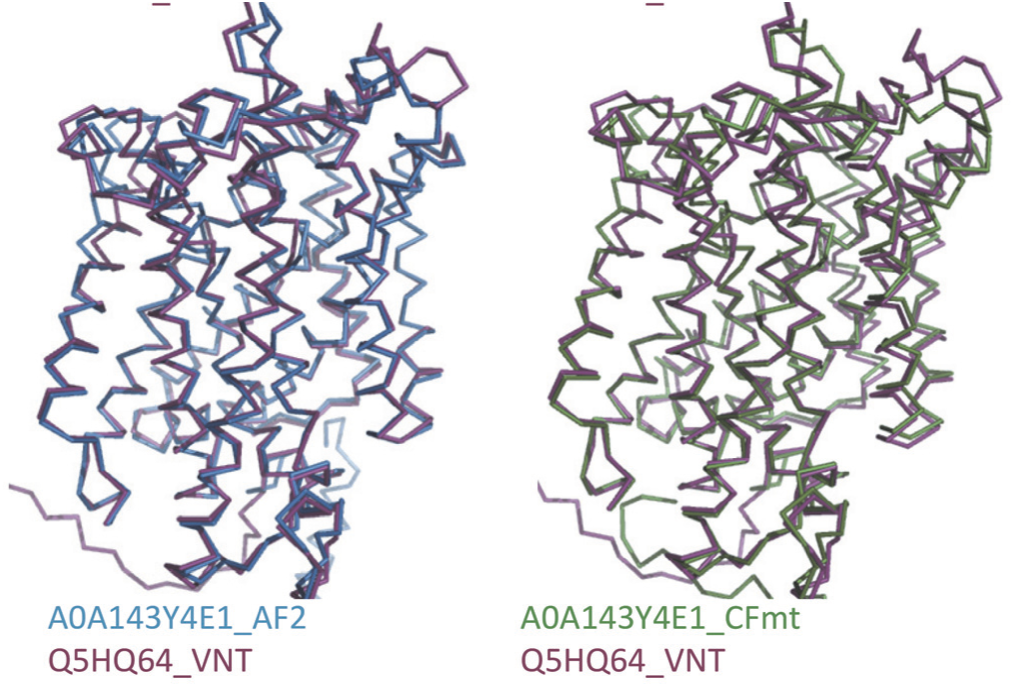
VNT compound mutagenesis of MCb_ie_ A0A143Y4E1 induces a conformational response like MCb_gut_ templates. **A**. Identical VNT mutations result in similar rearrangements of the models predicted by CFnt (MCb_ie_ A0A143Y4E1) or CFpdb (MCb_gut_ Q5HQ64), indicated by pRMSD from MCb_gut_ Q5HQ64 native CFpdb model (inward open). **B.** MCb_ie_ A0A143Y4E1 and MCb_gut_ Q5HQ64 VNT mutant 3D models are quasi-superimposable (top left).

However, detailing the effect of h3 YN and/or h6 VY ^+^/_-_ h6 T mutations showed that, unlike MCb_gut_, MCb_ie_ A0A143Y4E1 predicted conformation was fully switched in response to h3 YN mutation only (Figure 8AB), as previously observed using MCg1 A0A149PND7 [6]. Yet, contrary to MCg1 A0A149PND7, additional mutations targeting potential suppressor sites of YN-induced conformation switch had no impact on modeling of A0A143Y4E1 YN (Figure 8C).

**Figure 8.**
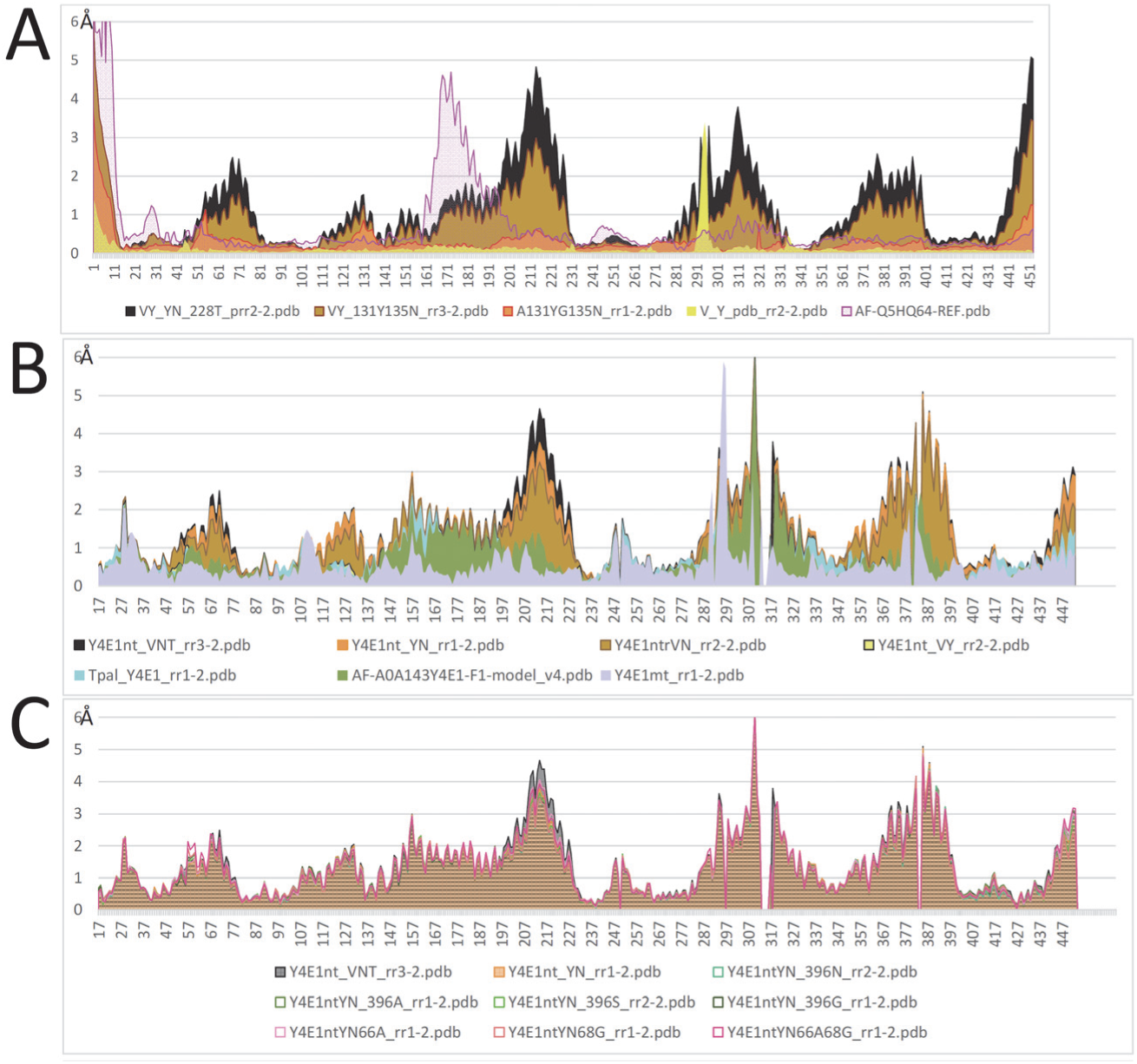
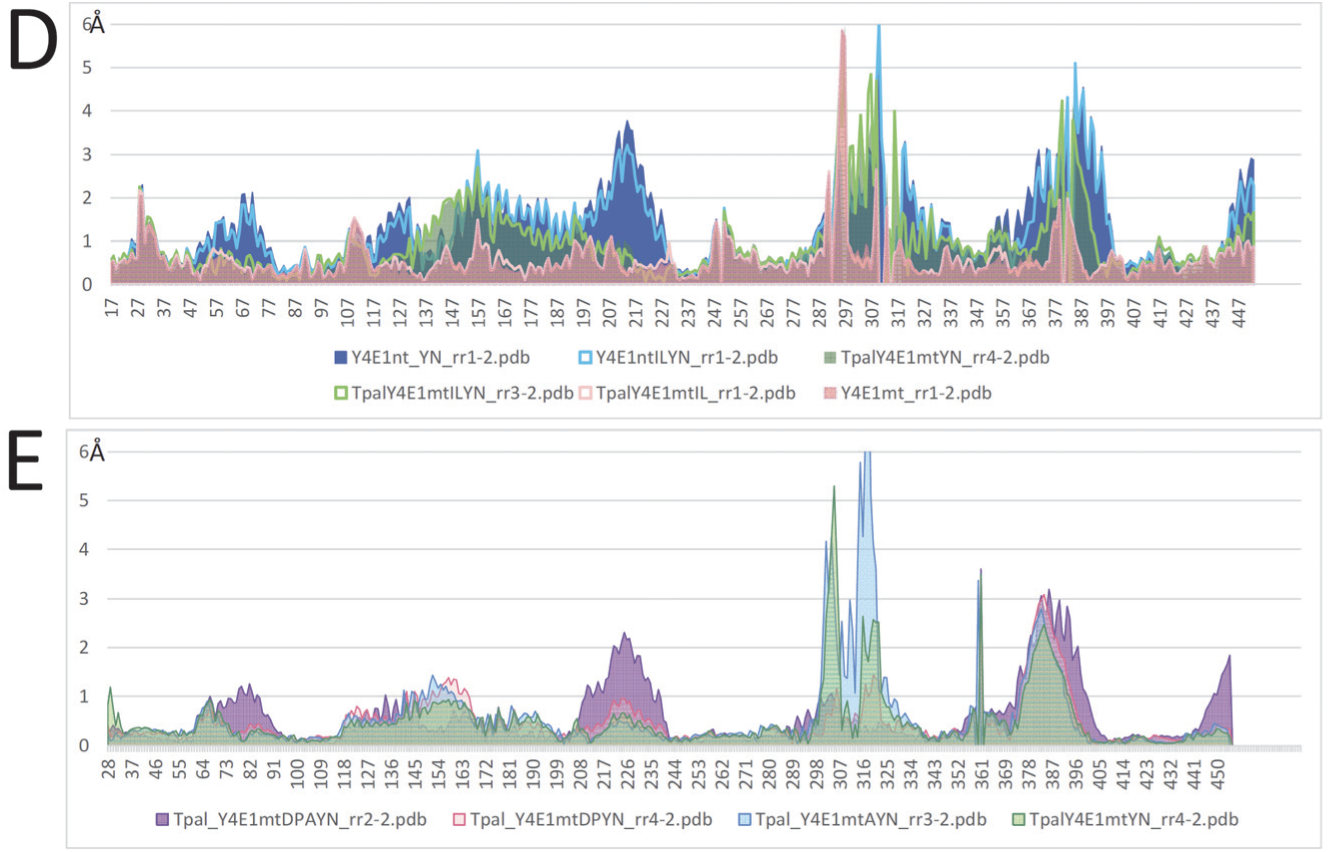
Distinct conformational response of MCb_ie_ A0A143Y4E1 to select mutations targeting sites of Slc11 synapomorphy. **A**. Recap of the effect of VNT mutagenesis on MCb_gut_ Q5HQ64 (h3 A131YG135N, h6 A228T M231V H233Y, [6]) shown as pRMSD from Q5HQ64 native CFpdb model (inward open). **B.** MCb_ie_ A0A143Y4E1 displays increased sensitivity to h3 mutation YN. **C.** Modeling of MCb_ie_ A0A143Y4E1 YN-induced conformation change is unaltered by secondary mutations previously shown to suppress MCg1 A0A149PND7 YN-induced rearranged modeling [6]. **D.** The impact of h3 YN mutation on MCb_ie_ A0A143Y4E1 CFnt modeling is lost in the compound mutant (A0A143Y4E1_15mt, cf Figure 5), and additional h3 mutations have no further effect. **E.** Reverting 3 of the 15 targeted mutations co-assayed in MCb_ie_ A0A143Y4E1_15mt restores some modeling responsiveness to h3 mutation YN.

In addition, A0A143Y4E1_15mt modeling was insensitive to h3 YN mutagenesis and neither A0A143Y4E1 YN, A0A143Y4E1_15mt YN nor A0A143Y4E1_15mt model were affected by adding 2 other h3 mutations (Figure 8D). Furthermore, reversing 3 coevolutionary rate-shifts that distinguish MCb_gut_ in l 7/8 (DPA sites #582, #589, #596 Figure 5A) restored a degree of conformational response to h3 YN mutagenesis, observed with A0A143Y4E1_15mt DPAYN model, akin to A0A143Y4E1 YN modeling response (Figure 8E).

These results indicate MCb_ie_ modeling operated the conformation switch toward the OO state in response to Slc11 synapomorphy VNT mutation, as expected from prior modeling studies of MCb_gut_ and MCg1. But MCb_ie_ A0A143Y4E1 modeling displays unique sensitivity to h3 YN mutation, which is lost after introducing 15 evolutionary coupled mutations that specify the clade MCb_gut_. These data support divergent structure/function relationships between MCb_gut_ and MCb_ie_ and suggest a conformational role for intervening loops that are distal from the Mn^2+^ binding site.

### 2.5. Conformation-dependent impact of mutations targeting MCb l 7/8 and l 8/9

Reversing MCb_ie_/MCb_gut_ rate-shifts in l 7/8 (DPA) was guided by observing they map to a specific area of MCb_gut_ structure (Figure 5A), i.e., not part of the evolutionary transition from MCa to MCb (Figure 3BC) and polymorphic (Figure S3). And preserving this area as is in MCb_ie_ retained some conformational flexibility in MCb_ie_ mutant A0A143Y4E1_15mt DPAYN (Figure 8E). Besides, the functional importance of these DPA residues for MCb_ie_ conformational response to h3 YN mutagenesis is underscored by their conservation in MCa clade (Figure S3, sites #582, #589, #596).

Along similar lines, mutation at site #632 (Figure 5A) in l 8/9 (Figure S3) was reevaluated. To mimic more precisely MCb_ie_/MCb_gut_ residue exchange, vs MCb_ie_/MCa exchange, both sites #632 and #633 (cf Figure S3) were initially mutated in A0A143Y4E1_15mt (counted as one mutation). To verify whether this dual mutation also introduced 3D constraint it was reverted as well, either alone or combined to l 7/8 DPA reversion mutation (Figure 9A). In both instances, restoring MCb_ie_ native sequence in l 8/9 recovered some conformational flexibility. Modeling of the amended mutant A0A143Y4E1_15mt DPA-KP responded to YN-induced conformation exchange like native A0A143Y4E1 (Figure 9C). However, in absence of the YN mutation, the amended mutant A0A143Y4E1_15mt DPA-KP deviated locally from reference MCb_gut_ Q5HQ64 IO conformer (Figure 9B) and did not appear superimposable anymore (Figure 9CD).

**Figure 9.**
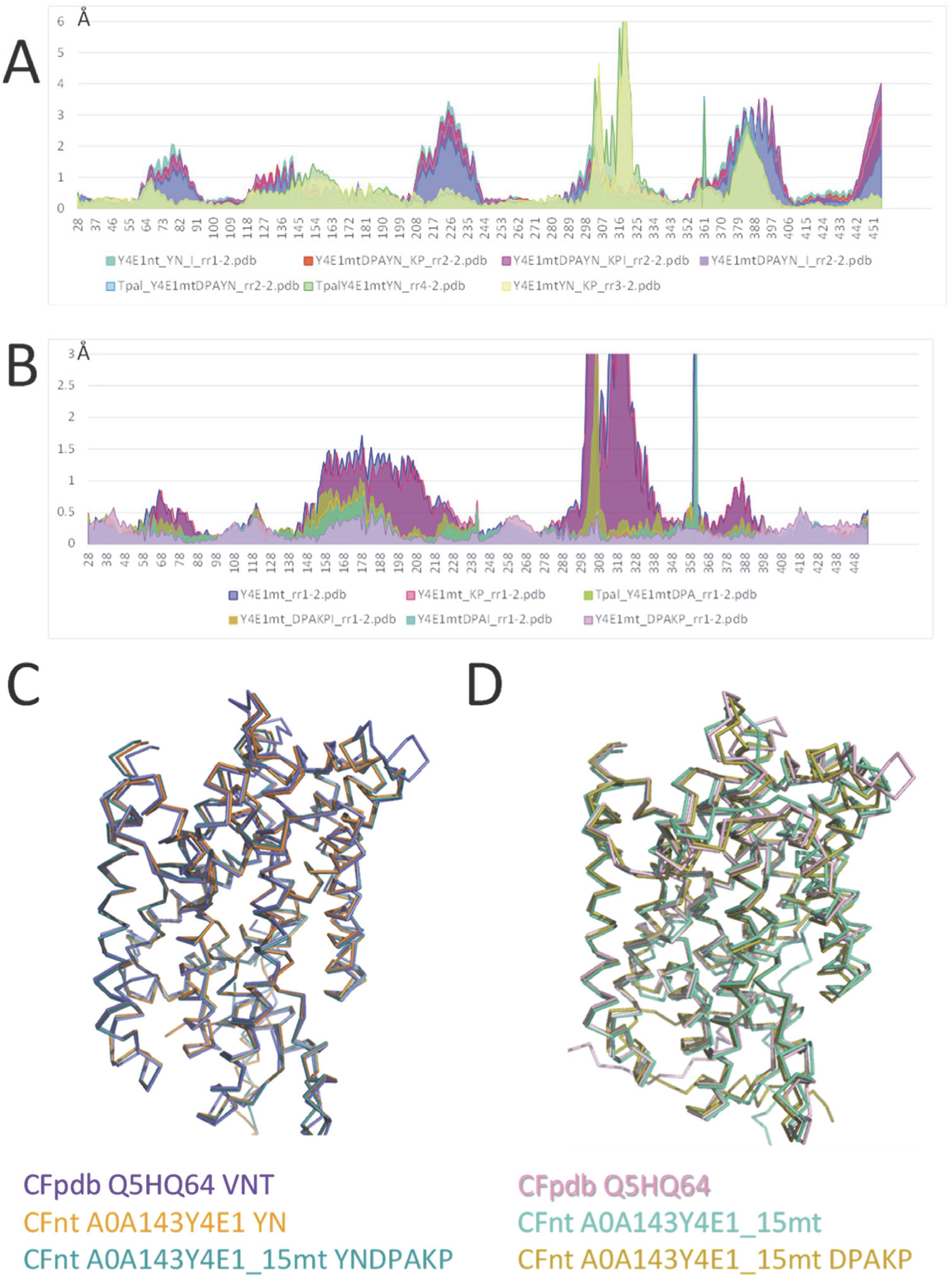
MCb_ie_ A0A143Y4E1 shows conformation-dependent responses to reversion of 5 of the 15 sites mutated in A0A143Y4E1_15mt. **A,B**. pRMSD relative to native A0A143Y4E1 CFnt model. **C,D.** Multiple 3D structure alignment using POSA. Reversion of 5 mutations from A0A143Y4E1_15mt has dual effects: it allows YN-induced conformation switch toward outward open state (**A, C**) but decreases 3D resemblance to MCb_gut_ Q5HQ64 inward open 3D model (**D**) by restoring instead similarity to the native MCb_ie_ A0A143Y4E1 model (**B**).

In sum, introducing 15 coevolutionary rate-shits that distinguish MCb_gut_ from MCb_ie_ into MCb_ie_ recipient A0A143Y4E1 allows modeling an IO conformer that is superimposable onto MCb_gut_ reference IO conformer (A0A143Y4E1_15mt). Conversely, reciprocal mutagenesis of MCb_gut_ members demonstrated native conformation-dependent effects, underscoring the functional significance of the targeted sites (cf section 2.3).

However, A0A143Y4E1_15mt lost 3D flexibility and responsiveness to YN-induced conformation exchange from IO state to OO state. This impact was alleviated by reverting 5 of the mutations that targeted extramembranous loops distal from the substrate binding site (l 7/8 & l 8/9). Yet these same reverted sites abrogated the 3D resemblance observed between the MCb_ie_ compound mutant and reference MCb_gut_ structure in IO state. These 5 sites therefore demonstrate conformation-dependent effect in the MCb_ie_ template A0A143Y4E1, and similar results were obtained using another MCb_ie_ template, A0A285P4W2 (from the Bacillaceae *Terribacillus aidingensis*, Figure S4).

To conclude, reciprocal mutations of coevolutionary rate-shifts that took place as MCb_ie_ and MCb_gut_ diverged exert distinct conformation-dependent effects on 3D modeling of members of each clade. This implies the targeted sites contribute to the mechanism of MCb carrier cycling albeit their respective role may differ within MCb_ie_ or MCb_gut_ model structure. Together these data relate MCb_gut_-specific functional divergence to the IO state of the carrier, which seems consistent with a possible shift in Mn import capacity in LB.

### 2.6. MCb_gut_ phylogeny reveals a pair of distinct carriers in the common ancestor of LB

To approximate a possible period for this functional transition, from marine MCb ^+^ Bacillaceae toward MCb ^+^ LB, MCb phylogeny was further explored. First, among spp. of the Enterococcaceae family because its origin was dated (>425 million years ago [20]) and its structure comprises four clades, one of which diverged basally [30]. MCb_gut_ phylogeny is complex due to pervasive horizontal gene transfers [31] and multiple copies per genome (e.g., 6 in *Levilactobacillus suantsaii* strain CBA3634 chromosome, NZ_CP059603.1). This may explain why the phylogenies of enterococcacal MCb_gut_ and Enterococci spp. vary (Figure S5). But the general topology of the tree and the profound dichotomy of *Enterococcus* clade A vs clades B-D were recovered, implying significant phylogenetic signal encoded by MCb_gut_. Separate grouping of MCb_gut_ seqs resulting from gene transfer to bacteria of different phyla, either enterobacterial or actinobacterial spp., indicate distinct events. The position of the Enterococcaceae *P. termitis* seq suggested it may demarcate 2 types of MCb_gut_, meaning that perhaps the common ancestor of Enterococcaceae possessed a pair of distinct *mntH* Cb_gut_ genes.

To address this possibility, MCb_gut_ phylogeny was determined across the LB order using a set of representative seqs from each LB family (Figure 10). The results show the presence of two clusters of MCb_gut_, labeled MCb_gu1_ and MCb_gu2_, found both in Enterococcaceae and Lactobacillaceae. In addition, the data suggest two other clades could diverge independently from MCb_gu2_.

**Figure 10.**
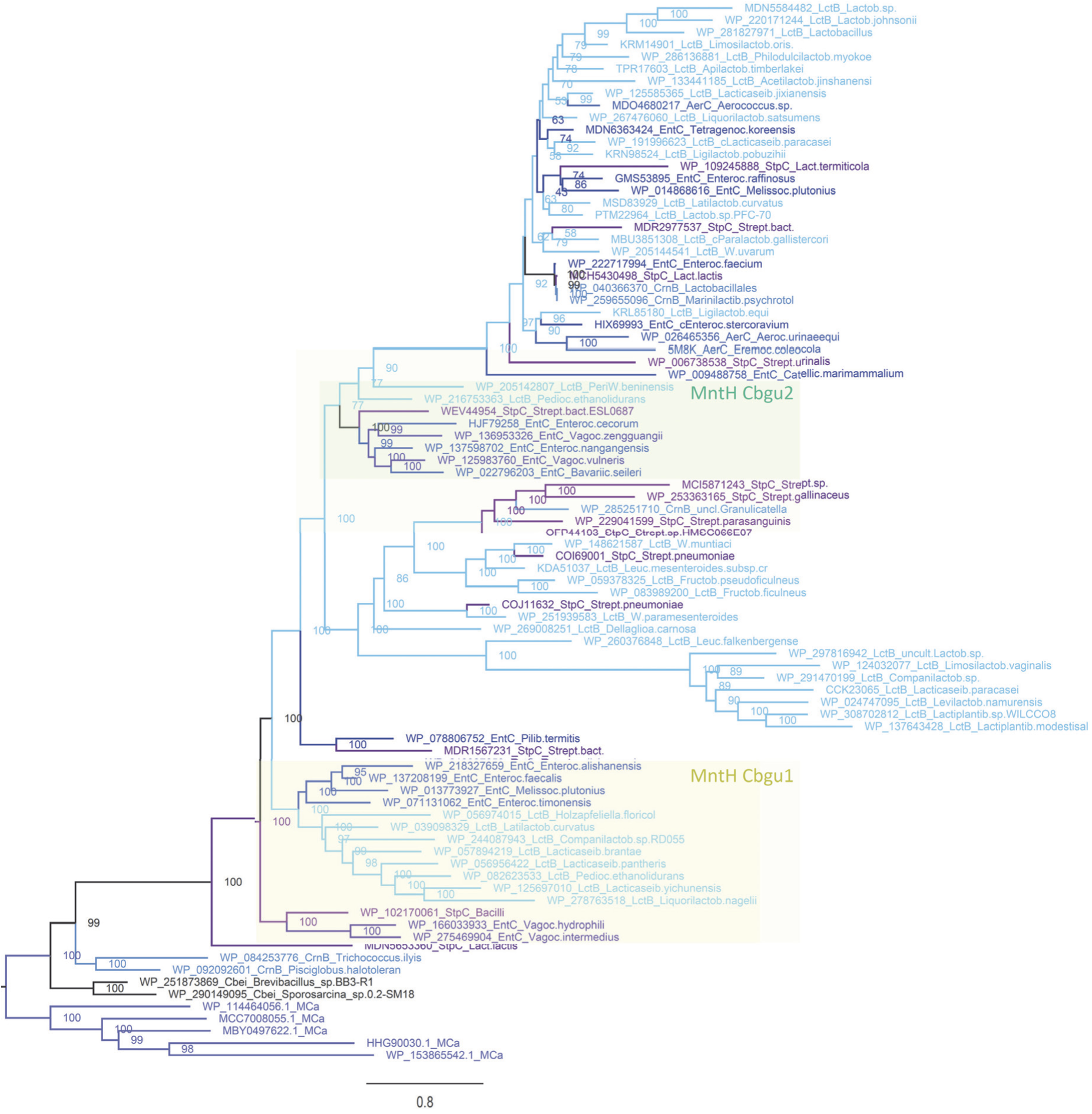
IQ-Tree phylogeny of MCb_gut_ among Lactobacillales (LB). Representative seqs from each LB family (Aerococcaceae, AerC, Lactobacillaceae, LctB, Streptococcaceae, StpC, Enterococcaceae, EntC, Carnobacteriaceae, CrnB) were analyzed using 5 MCa and 2 MCb_ie_ seqs as outgroup. The tree presented used 347 PI sites, the substitution model EX-EHO, ML estimate of a.a. state frequency, free rate model of variation among sites with 7 categories. The statistical significance of each node is indicated together with the scale (nb substitution per site). Two clusters of MCb_gut_ seqs, separated by *P. termitis* seq and that replicate the profound dichotomy observed by analyzing MCb_gut_ seqs from Enterococcaceae (Figure S5) are highlighted: MCb_gu1_ and MCb_gu2_.

The present tree thus supports the possible existence of two MCb_gut_ copies in the common ancestor of LB. Several spp. of Lactobacillaceae harbor both genes encoding MCb_gu1_ and MCb_gu2_ on their chromosome (e.g., NZ_LT854705.1, NZ_CP059603.1) implying perhaps some LB retained both genes since their origin.

The presence of both MCb_gu1_ and MCb_gu2_ in the common ancestor of LB suggests they are not functionally equivalent. To evaluate this proposal 3 individual sequences from each type were subjected to AF/CF modeling using either native sequences or compound mutants. Native AF2 models showed that, both in Lactobacillaceae and Enterococcaceae, MCb_gu1_ and MCb_gu2_ yielded distinct conformers in the OO state (Figure 11A & 11B, respectively). MCb_gu2_ type appeared closer to the fully OO conformation and less structurally conserved among Enterococcaceae. Introduction of the set of (reciprocal) mutations corresponding to MCb_ie_/MCb_gut_ evolutionary transition (cf section 2.3) confirmed it had little impact on OO conformers of MCb_gu1_ and MCb_gu2_ types (not shown). These data are in accordance with previous observations showing a strong phylogenetic component in AF/CF modeling of MCb_gut_ [6].

**Figure 11.**
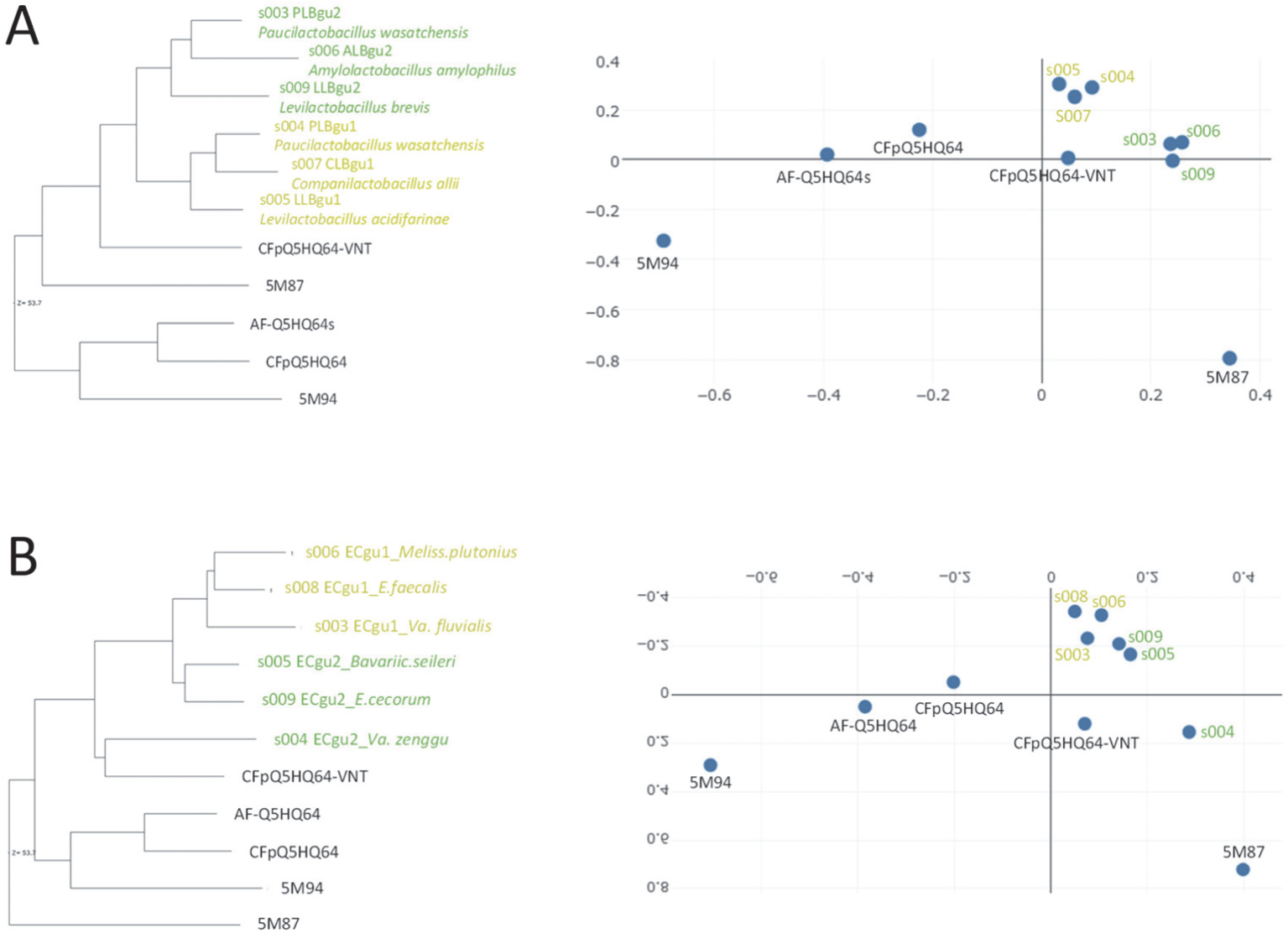
AF2 models distinct conformers for LB MCb_gu1_ and MCb_gu2_ templates. **A**. Lactobacillaceae. **B.** Enterococcaceae. Pairwise structural similarity was established with Dali all against all using as 3D references the solved structures of MCb_gut_ in outward open state (5M87) and inward open state (5M94) as well as AF2 and CFpdb models of MCb_gut_ Q5HQ64 and CFpdb model of Q5HQ64 VNT. Similarity depicted using dendrogram (left) and 3D correspondence (right) representations.

To compare MCb_gu1_ and MCb_gu2_ structural dynamics a combination of mutations able to mimic carrier forward transition, i.e. exchanging the OO state modeled by CF pdb using the native sequence for the IO state of a mutated carrier, was necessary. Along sites previously identified as conformationally active, either part of Slc11 synapomorphy or contributing to inner gate opening, additional sites were targeted in helices 3, 4, 8 and 9 which together constitute Slc11 carrier ‘hash module’ known to interact with the pmf [2], [3]. All targeted sites are evolutionary conserved so that identical mutations were introduced among both MCb_gu1_ and MCb_gu2_ templates (Table S2).

Introducing a series of 15 phylogenetically aware mutations into the *Paucilactobacillus wasatchensis* template PLB_gu1_ enabled modeling an IO conformer like MCb_gut_ Q5HQ64 reference CF model (Figure 12AC). Applying this combination of mutations to other MCb_gu1_ had similar yet less pronounced effect, producing models more advanced toward, albeit not switched to, the IO state (Figure 12E and Figure S6A,C). In comparison, MCb_gu2_ mutant response appeared more sequence-specific with two templates (Ece_gu2_, Lbr_gu2_) progressing toward the IO state (Figure 12F and Figure S6B,D).

**Figure 12.**
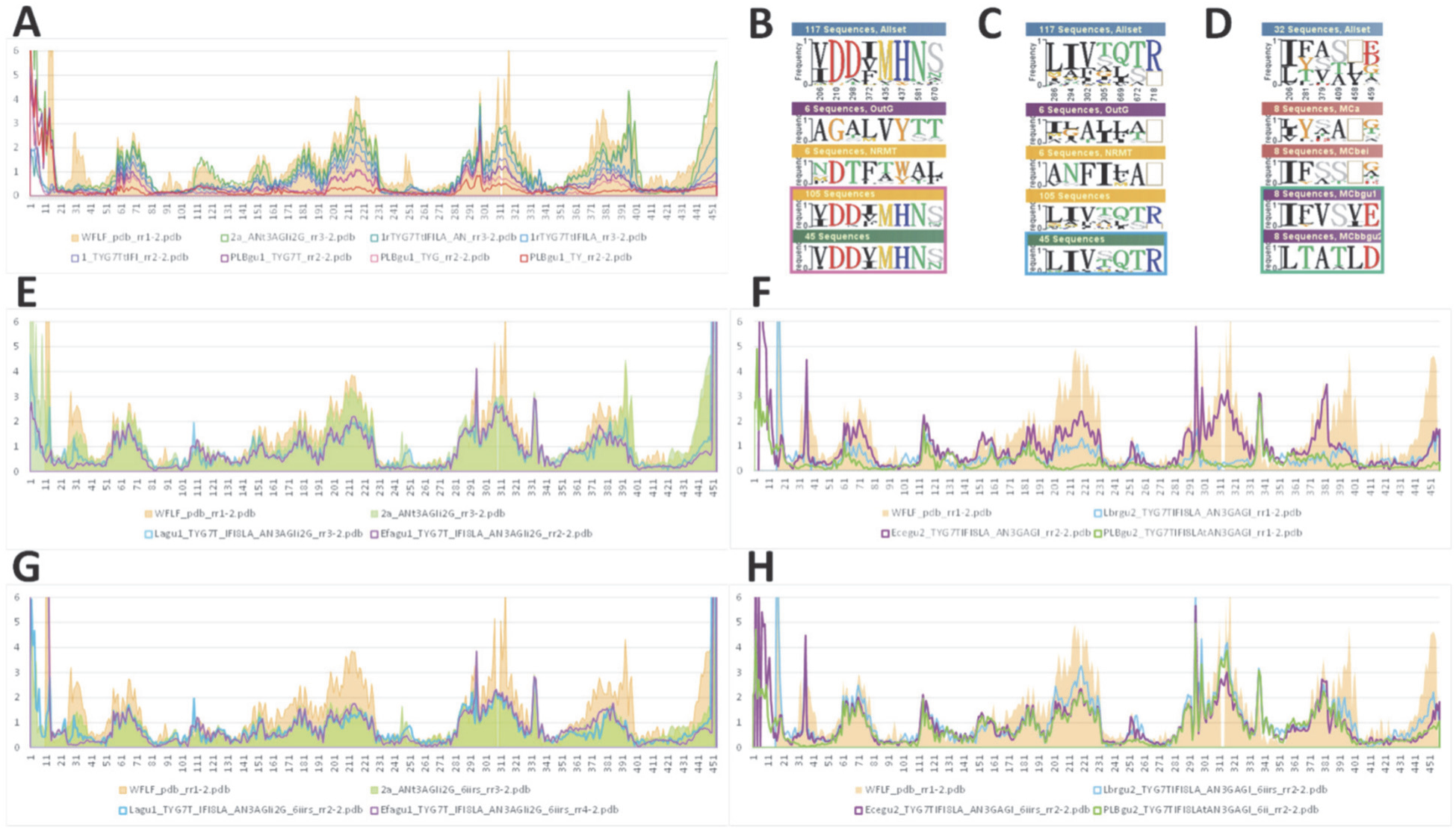
CFpdb modeling reveals functional differences between MCb_gu1_ and MCb_gu2_. **A, B, C**. Mutagenesis driven conformational change, from the outward open state toward inwardly open state, obtained by CFpdb modeling of compound PLB_gu1_ mutants is shown as pRMSD from native PLB_gu1_ model. The a.a. replacements targeted selectively conserved sites either across the Slc11 family (**B**) or among prototype Nramps (pN-I, pN-II) and MntH C clades (MCa, MCb, MCg and MCaU, [7]) (**C**). **E, F.** Identical compound mutants LLB_gu1_ (Lagu1) and EC_gu1_ (Efagu1) display conformational responses like PLB_gu1_ while the mutants PLB_gu2_, LLB_gu2_ (Lbrgu2) and EC_gu2_ (Ecegu2) exhibit more diverse overall shapes, shown as pRMSD from EC_gu1_ (**E**) and PLB_gu2_ (**F**) native 3D models. **D, G, H.** Adding reciprocal mutations to exchange residues that distinguish MCb_gu1_ from MCb_gu2_ (**D**) impaired MCb_gu1_ conformation similarly for all 3 templates (**G**), while some MCb_gu2_ showed opposite effect (**H**), indicated by pRMSD from EC_gu1_ (G) and PLB_gu2_ (H) respective native 3D models.

To further distinguish the modeling properties of MCb_gu1_ and MCb_gu2_ templates a set of discriminatory type ii evolutionary rate shifts was targeted to perform reciprocal residue exchanges between MCb_gu1_ and MCb_gu2_ seqs (Figure 12D). Again, all 3 MCb_gu1_ behaved similarly and were modeled into more OO conformers, implying some antagonism between the two series of mutations tested altogether (Figure 12G and Figure S6E). Reciprocal mutagenesis had distinct effects on MCb_gu2_ templates, which appeared either unchanged or further progressing toward the IO state (Figure 12H and Figure S6F).

Lastly, evolutionary conservation among representative sets of MCb_gu1_ and MCb_gu2_ seqs suggests these clades experienced distinct selective pressures (Figure 12 I&J, Figure S7).

From these results it is concluded MCb_gut_ phylogeny replicates the evolutionary dichotomy previously established among *Enteroccoccus* spp. It appeared the common ancestor of LB likely possessed a pair of genes encoding seemingly distinct types of MCb_gut_. Transition from marine Bacillaceae MCb_ie_ toward LB’s MCb_gut_ could proceed sequentially with functional evolution of a substrate-dependent step, such as inner gate opening, followed by gene copy and separate evolution of distinct carriers.

## 3- Discussion

This in silico study correlates the emergence of LB with the functional divergence of a pair of (Slc11) MCb_gut_ carriers. MCb_gut_ carriers share a common ancestor with their sister clade MCb_ie_, and together they form the group MCb which evolved from the group MCa. MCb_ie_ predominates in marine Bacillaceae and demonstrates evolutionary conservation; it relates directly MCa to MCb_gut_ because MCb_ie_ seqs display non overlapping patterns of identical residues with relatives from each group. Comparative studies between the sister clades MCb_ie_ and MCb_gut_ indicate functional innovation to open the inner gate of MCb_gut_ carriers as well as MCb_gut_-specific structural dynamics plasticity. Apparently, MCb_gut_ could evolve from MCb_ie_ in response to selective pressure that fostered functional divergence. Assuming this was the case, such evolutionary pressure indicated novel environmental conditions. It is hypothesized that the gastro-intestinal tract of early marine vertebrates provided a novel niche conducive for progressive evolution of a MCb ^+^ Bacillaceae into the ancestor of LB, which carried a pair of genes coding for distinct MCb_gut_.

To what extent the data gathered support the possibility that Mn-dependent, MCb ^+^ LB evolved from some Bacillaceae precursor adapted to marine environment? First, several bacterial orders enriched in MCa^+^ spp. (e.g., Cyanobacteria, Pseudomonadales (gamma proteobacteria, GPB), Burkholderiales (BPB), Rhodobacterales, Sphingomonadales, Hyphomicrobiales (Rhizobiales, APB) [7] might have formed Precambrian subaerial microbial mats that could include *Bacillus* spp. [32], [33]. Such a terrestrial setting would facilitate the acquisition of *mntH Cb_ie_*gene precursor (*mntH* Ca) by some *Bacillus* spp. Bacillaceae represent the primary source of *mntH Cb_ie_* gene, consistent with evidence of recent gene transfers toward different spp. of Paenibacillaceae (*Paenibacillus* and *Brevibacillus* spp.) in which *mntH* Ca dominates. And many MCb ^+^ genera are marine Bacillales found in deep-sea sediments (e.g. *Bacillus*, *Paenibacillus*, *Lysinibacillus* and *Terribacillus* spp.) while others populate various environments, including the gut of animals [34], [35].

Second, while MntH A (MA) type, which is ancestral to all *mntH* C genes that were derived from eukaryotic cells (Figure 1A, Table S1), predominates in Bacillaceae, a fraction of Bacillaceae spp. acquainted with marine environment adopted instead MCb_ie_. MCb_ie_ phylogeny indicates genera such as *Lederbergia*, *Siminovitchia*, *Heyndrickxia*, and *Neobacillus* (Figure S1B, [36], [37]) may have inherited *mntH* Cb_ie_ vertically. Their phylogeny points these species as more likely candidate precursors of LB than MA^+^ Bacillaceae, such as *B. subtilis* and *B. cereus*. Also, these marine Bacillaceae grow aerobically, which implies Fe-based metabolism, and some were recovered as well from dairy environment or found in the gut of various animals. Emergence of MCb_ie_ from MCa in marine Bacillaceae could thus mark a first step in the adaptation to the gut of animals such as invertebrates.

A consolidated hypothesis may thus be viewed in successive steps (Figure 13) with i) emergence of MCa^+^ Bacillaceae within subaerial/terrestrial microbial mats, which also colonized marine habitat, ii) adaptation of marine MCb ^+^ Bacillaceae to the gut of invertebrates, and iii) selection of Mn-dependent LB carrying a pair of *mntH* Cb_gut_ genes in the gut of early vertebrates. Previous studies suggest exchange of MA for MCa affected the coupling of the pmf to Mn^2+^ import [7], which could improve resistance to environmental stresses (e.g., dessication, temperature, osmotic stress) and provide a path toward adaption to acidic environment through evolution of MCa to MCb_ie_, and then to MCb_gut_ – a step shown to impact opening of the inner gate to carry Mn^2+^ ions inside cells. Prior works on MCb_gut_ homologs revealed both the plasticity of their functional dynamics [6] and their efficient use by LB to pump Mn^2+^ in competitive environments [12].

**Figure 12.**
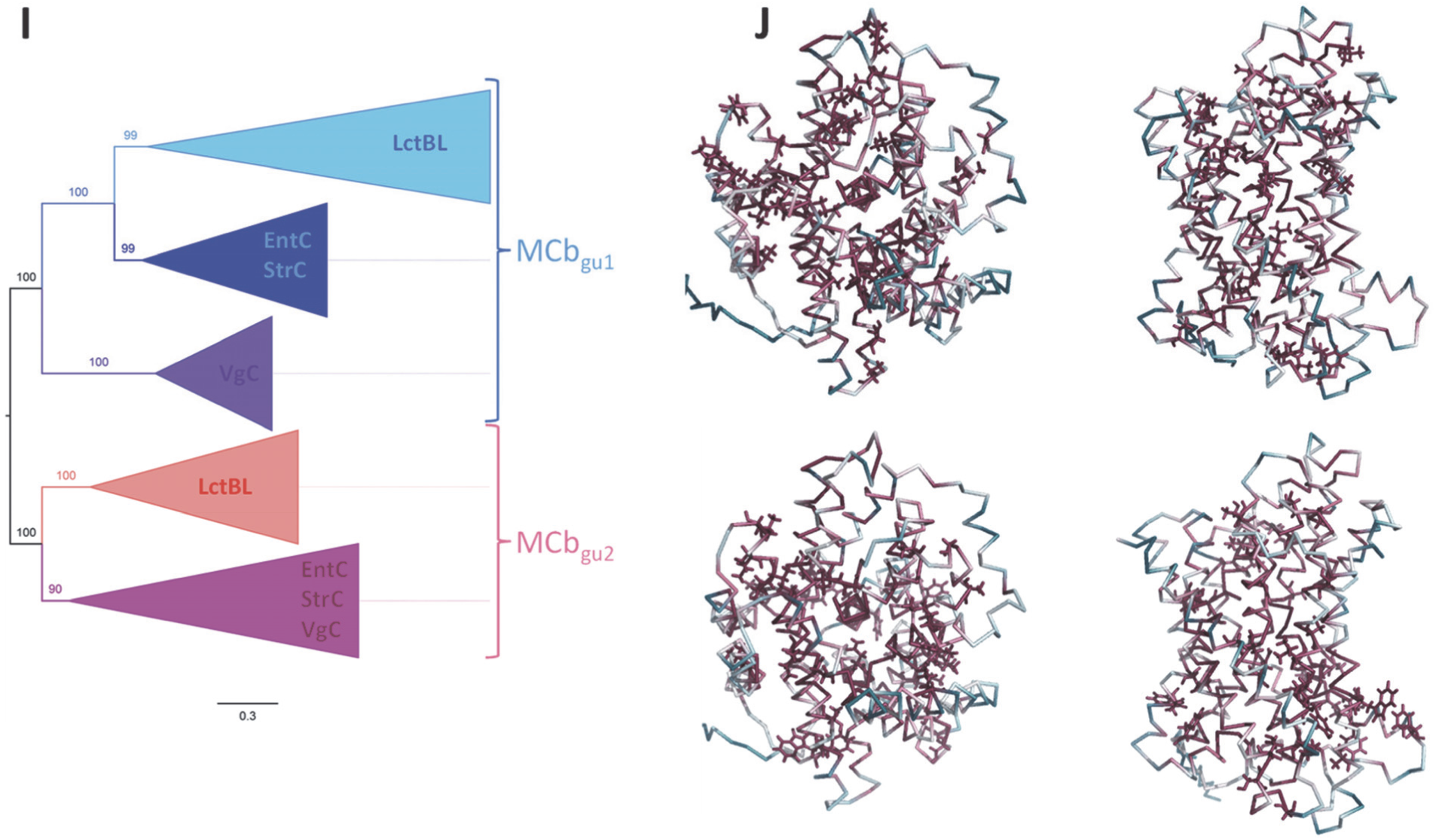
ctn’d**. I, J.** Evolutionary divergence of MCb_gu1_ and MCb_gu2_ clades. Simplified IQ-Tree representation of MCb_gu1_ and MCb_gu2_ clade relationships (I, 46 seqs each, detailed in Figure S7). Clade-specific Consurf evolutionary conservation patterns among MCb_gu1_ (top, native EC_gu1_ CF model, Efagu1) and MCb_gu2_ (bottom, native EC_gu2_ CF model, Ecegu2) clades (J). Model MCb carrier views of the outward face (left) and the transmembrane helix bundle (right).

**Figure 14.**
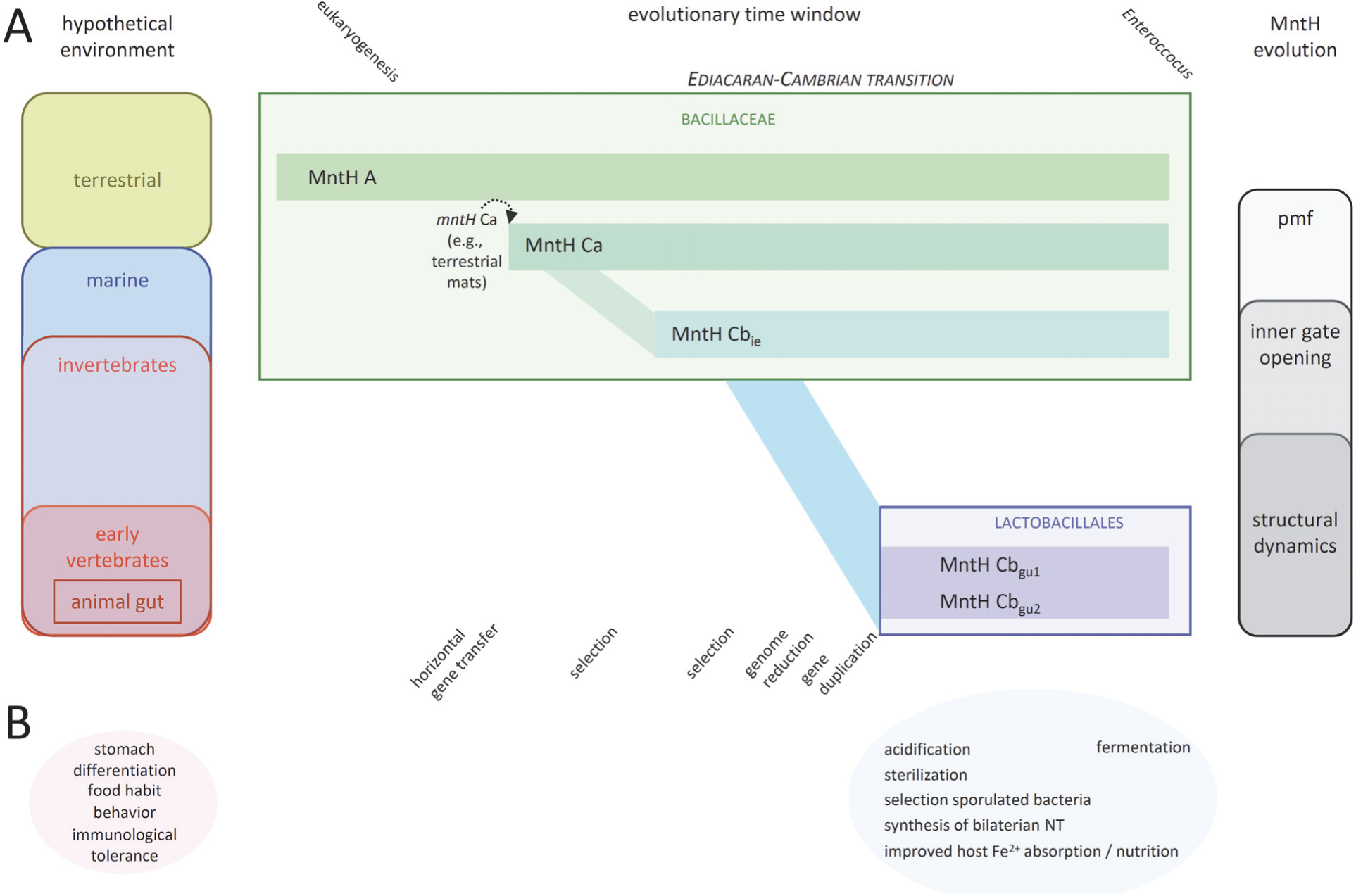
Hypothetical emergence of LB in the gut of early vertebrates based on molecular genetic and structural evolutionary analyses of Slc11 clade MCb (MCb_ie_ and MCb_gut_, i.e., MCb_gu1_ and MCb_gu2_). **A.** The evolutionary time window considered is indicated, from eukaryogenesis, ca. 2 billion years ago, to emergence of the genus *Enterococcus*, ca. 425 million years ago. It is proposed LB evolved during the Ediacaran-Cambrian transition (635-485 mya), possibly in the early Cambrian (ca. 540 mya). Substitution of the ancestral *mntH* A gene for a horizontally acquired *mntH* Ca gene is posited as initial step (MntH Ca^+^ Bacillaceae) followed by a combination of selective processes, including genome reduction and gene duplication. Successive environmental changes that could favor evolution of MCb (left) and previously reported stepwise changes in MntH functional properties (right) are indicated (from top to bottom). **B.** LB colonization of the gut of protochordates led to acidify the upper part of the gut during food fermentation, producing outcomes that could foster the establishment of the vertebrate gastro-intestinal tract in its canonical form (right panel), by emulating developmental host functions such as differentiation of the stomachal pouch, food habits and behavioral responses, and immunological tolerance (left panel).

Whether this LB evolution toward Mn based metabolism matched the final design of the gastro-intestinal (GI) tract of early vertebrates is a matter of conjecture. But this possibility may provide an interesting angle to approach our intimate relationships with LB because vertebrate development owed to successive innovations following differentiation of the pharynx at the entry of the digestive tract in chordates, including for instance the development of sensory organs and vertebrate novelties such as the spleen, individualized pancreas, heart innervation and eye’s own muscles [38], [39], which might have fostered mutualistic interactions between emerging LB and their host.

Vertebrate inventions took place around the 1R whole genome duplication (WGD) that occurred in the early Cambrian and which produced, in particular, the archetype Nramp ‘Ohnologs’ Nramp1 and Dmt1 ([40], [41], [42] and Figure S8). *NRAMP1* (*SLC11A1*) expression depends on C/EBP [43], [44] and dominates in the spleen, blood and tissue phagocytes, contributing to host resistance to infection [8], [45]. Dmt1/Nramp2 (Slc11a2) catalyzes H^+^-dependent Fe^2+^ import both as part of the transferrin endocytic cycle that delivers iron through the body and at the apical membrane of duodenal enterocytes [46], [47]. This context may suggest the Ediacaran transition to early Cambrian as possible evolutionary interval for LB emergence while the GI tractus of early vertebrates was developing into its final form.

High efficiency Mn intracellular accumulation under acidic conditions makes physiological sense for LB because Mn as a micronutrient accelerates their growth under lactate stress [48]. Lactic acidification of LB environment has important effects on the GI tract: inhibiting the growth of potential microbial pathogens while providing a nutrient for potentially beneficial bacteria, such as spore-forming component of the colonic microbiota [49], which by consuming lactate that is transformed into short chain fatty acids (SCFA) may stimulate LB growth in return [50], [51]. These effects, added to LB ability to predigest complex foods could possibly help develop the GI tract of early vertebrates by favoring differentiation of the gastric organ (stomach).

The GI tracts of urochordates and cephalochordates comprise organs homologous to the liver and pancreas of vertebrates, and chordates share the molecular basis of extracellular digestion with vertebrates. Yet, their genomes lack the *pepsinogen* gene encoding a gastric digestive enzyme and cephalochordates demonstrate evidence of intracellular digestion in the hepatic cecum. This suggests that extracellular digestion systems based on both stomach and pancreatic digestive enzymes emerged specifically in the common ancestor of vertebrates ([52], [53] and based on homologs of GTEx stomach-specific encoded polypeptides, [54], detected in Cephalochordates: Lipase F, gastric type >50% aa id; in Chondrichthyes: Pepsinogen >60% aa id; in Tetrapods: Trefoil factor 1 & 2, >67% aa id).

LB strains isolated from mammalian source (mainly rodents) colonize the gastric epithelium by forming biofilms [18, 55], [56], [57]. To resist the harsh environment of the stomach lumen LB use glutamate and aromatic L-amino acid decarboxylases which help sustaining the transmembrane pmf to energize metabolism and transport [58], [59]. The fact these bacterial enzymes produce amino acid derived compounds used as neurotransmitters (NT) in bilaterian animals [60], such as serotonine, dopamine and GABA [56], both in the central nervous system (CNS) and the enteric nervous system (ENS) [61], suggests a possible feed forward loop [62] where proto-LB residing in the upper gut of some vertebrate ancestor would respond to food intake by generating an acidic environment facilitating digestion and contributing to gut-brain signaling.

An acidic environment in the duodenum of vertebrates is key for absorbing the growth supporting micronutrient iron in the form of Fe^2+^ [47]. As it seems, selection of duodenal Slc11a2– and H^+^-dependent Fe^2+^ uptake may thus correspond to the period of establishment of the vertebrate stomach. And the prospect these processes were supported by Mn^2+^-dependent acid-resistant LB expressing MCb_gut_ parologs may bear significance in terms of distinct substrate selectivity among Slc11 homologs, and partition of micronutrient resources for mutual benefits [63], to the vertebrate host (Fe) and to the LB residing in its gut (Mn). Arguably, an example of ecological-evolutionary interaction [62], [64], [65] that provided a rich foundation to diversify vertebrate holobionts.

Previous in silico data obtained with MCb_gut_ demonstrated highly plastic structural dynamics that may increase Mn import efficiency, hereby sustaining LB resistance to acidic and osmotic stress. Herein, further study of MCb_gut_ revealed a parology that predated LB radiation while the sister carrier MCb_ie_ showed structural differences likely altering Mn intracellular release. These data may suggest evolution of LB MCb_gut_ could foster a novel, Mn-dependent way of life in an acidic environment stock up with food while relinquishing use of Fe to their host. This process was likely progressive, contracting LB’s ancestral MCb_ie_^+^ Bacillaceae genome, and going along with differentiation of the stomachal pouch, as specialized organ in the neo gastro-intestinal tract of the ancestor of jawed vertebrates.

Gastric LB could impact host behavior and food habits through synthesis of NT that act directly on leukocytes within the GI tract lamina propria and affect nerve ganglia of the ENS, producing signals relayed through afferent vagal connections to the CNS [66], [67]. These instructions could be modulated by intestinal SCFA producing bacteria, which may act directly on vagal intestinal terminals and stimulate NT signals [68]. In return, efferent vagal influx from the CNS toward the spleen produces NT that directly regulate the secretion of proinflammatory cytokines by leukocytes [69]. Similarly, enteric bacteria could also modulate local steroid secretion, contributing to reduce stress hormone levels and inform the CNS to adjust behavioral output [61]. It would seem possible that emergence of LB owed at least partly to their ability to communicate with their host via the gut-brain-axis.

Vertebrate transition to extracellular intestinal digestion presented inherent risks that required immunological surveillance, because luminal microbes opting for a pathogenic lifestyle may weaken the intestinal barrier integrity and create a portal for host infection [70], [71]. Viewing regulatory lymphocytes as metabolic sensors, Treg mediating suppressive functions when food is scarce whereas inflammatory functions of Th17 are fueled by glycolysis [72], [73], suggests food digestion potentially incurs intestinal inflammation that depends on subsequent resolution to maintain homeostasis and develop immune tolerance to food-derived determinants [74]. Also, lactic acid sensing has anti-inflammatory effect by modulating macrophage phenotype polarization [75], [76]. According to these views, gut LB and SCFA producers and their metabolites could directly instruct the emerging adaptive immune system while sustaining systemic innate immunity, due to Firmicutes intrinsic lysozyme sensitivity [77].

Also, both lymphocyte and erythrocyte cell types emerged as vertebrate-specificities [78]. Duodenal Fe absorption boosted with gastric acidification likely helped developing the erythron (e.g., hetero-tetrameric hemoglobin in jawed vertebrates, [79]) to distribute oxygen within tissues and sustain their metabolic activity. Apparently thus, LB Mn-based metabolism and gastric acidification could enhance host Fe– and oxygen-based metabolism. And combined to their behavioral and immunological influence, LB role in stimulating host metabolism could elevate energetic efficiency and help diversify early vertebrates. Divergence of LB families may have been contemporaneous, perhaps reflecting various mutualist interactions implemented by diverse hosts.

According to recent views, the Ediacaran-Cambrian transition, which produced an animal radiation, could owe to a combination of causes, including transient catastrophic events (snowball earths, hypoxic environments) and progressive ‘biotic replacement’ through successive pulses of evolutionary innovations and ecosystem engineering [80], [81], [82], [83], [84], [85], [86]. The 1R genome duplication that occurred prior to divergence of jawed vertebrates may indeed testify of disruptive environmental changes during this period. On the other hand, the ability of extant LB probiotics to regulate mammalian brain, digestive and immune activity [56], [61], [67] suggests that emerging Mn-dependent MCb ^+^ LB colonizers, adapted to the upper gut of ancestral vertebrates, could impact the feeding habits and behavior of their host thereby influencing its evolution [87], [88].

Hence a LB-driven feed-forward loop may have contributed to diversify early vertebrate body plan, and turning to jawless vertebrates may be instructive in this regard. The sea lamprey has a life cycle comprising 3 distinct stages: blind larvae burrowing into the sand feeding on microbes and detritus by filtering surrounding water; a first metamorphosis rearranges sensory organs, forms a new esophagus and modifies the digestive system of free living juvenile lampreys that display high diet conversion efficiency and temperature-dependent growth; a second metamorphosis atrophies the intestine of fasting adults that die after spawning [89], [90], [91]. Apparently, the microbiota associated with growing wild juveniles is selectively enriched in LB, *Streptococcus* spp., in particular. Regarding hagfish gut microbiota, a close relative of LB *Weissella cibaria* (MCb ^+^) was detected in wild *Eptatretus* while in captive specimens, MntH A^+^ enterobacteria dominate (Figure S9). These data support the notion that gut LB could directly contribute to the growth and development of early vertebrates. However, continuous host development, perhaps tributary of further innovations such as jaws, heterodimeric hemoglobin and myelinization as well as the 2R genome allotetraploidization [40], [41], [79], [92] was likely required to further vertebrate’s radiation and inflate marine biodiversity by fuelling an evolutionary arms race toward ever more efficient predators [82], [93].

In sum, it is proposed that MCb ^+^ LB emerged in the upper gut of early vertebrates (ca. 540 mya, i.e, prior to vertebrate 1R WGD). LB colonization formed a novel holobiont that prefigured the final design of the gastro-intestinal tract of jawed vertebrates, whose development could benefit from boosted nutritional efficiency procured through LB-host interactions. Establishing a feed forward loop for food access and tolerance could incentivize LB genome reduction and adaptation as Mn-dependent gastric aids while their pro-gastric functions stimulated vertebrate Fe-dependent growth and development. Accordingly, LB evolution involved successive phases: i) reductive, during marine MCb ^+^ Bacillaceae adaptation to early vertebrate upper gut leading to Mn-centric, acid-resistant fermentative cells (MCb ^+^ LB); ii) expansive, through subsequent dissemination of MCb ^+^ LB adapting to novel niches, including terrestrial environments. Molecular communication between extant LB and mammalian gut brain-axis and immune system may include remnants of early vertebrate pro-gastric functions and subsequent interactions between more evolved organisms.

## 4- Materials & Methods

Protein sequences were collected using NCBI and Uniprot repositories and BLAST programs (PHI-BLAST, TBLASTN and BLASTP, [94]). Sequences were curated using the MPI-toolkit and the MMseqs2 program to retrieve sets of diverse seqs with reduced complexity [95]. ClustalX [96] seqs clustering and graphic display was used to unambiguously classify seqs into established phylogroups, using reference seqs [7].

Each phylogenetic analysis was performed using a balanced set of groups of seqs to be analyzed plus a defined outgroup. Seqs were multiply aligned using Muscle [97] and the resulting alignments were manually edited using Seaview [98]. As needed, Phylo-mLogo [99] display of the multiple sequence alignment was generated by writing a simple tree, aggregating clade-specific sets of seqs. Multiple sequence alignments were trimmed to eliminate sites with gaps or judged poorly informative in Phylo-mLogo display, using Treecon [100]. Parsimony-informative sites were selected using Mega [101]. Tree calculations with IQ-Tree [102] used a number of categories of free-rate heterogeneity model that was commensurate with the length of the parsimony-informative site alignment to estimate variations of substitution rate among sites using proportions of site categories and mixture substitution matrices [103]. Trees were displayed using FigTree (http://tree.bio.ed.ac.uk/software/figtree/) to appreciate the statistical robustness of tree nodes.

Site selection for mutagenesis used defined phylogroups of aligned seqs to identify candidate type ii evolutionary rate-shifts distinguishing the groups under study [4] based on the Sequence Harmony and multi-Relief approaches [104]. Sites producing highly significant scores with both approaches were selected as sets of coevolved sites discriminating sister phylogroups. Reciprocal mutagenesis was performed by exchanging sets of phylogroup-specific residues between templates from each group. Additional mutations targeted sites defining the family of Slc11 carriers (Slc11 synapomorphy) as well as sites that characterize clades part of the Slc11 family. Every point mutation introduced was verified a posteriori with the returned seq of the predicted model. Differences in evolutionary rates between sister clades were also visualized using Consurf [105].

3D modeling was performed using CF [11] with or without template (CFpdb and CFnt, respectively) and native seqs to identify the training model producing the best model, which was used to analyze corresponding mutant seqs [6]. AF2 models were retrieved from Uniprot repository [106]. Molecular analyses using 2StrucCompare [107] allowed to determine per residue root mean square deviation of the Ca trace between pairs of aligned structures. The All against all program (Dali EBI, [108]) was used to establish the 3D correspondence of groups of structures based on pairwise comparisons. POSA [109] was used to obtain multiple flexibly superimposed structures. Structures were visualized using Pymol (DeLano,W.L. The PyMOL Molecular Graphics System; Delano Scientific: San Carlos, CA, USA, 2002).

## Supplementary Materials

The following supporting information can be downloaded at

Figure S1. MCb (MCb_ie_ and MCb_gut_) phylogeny among bacterial families comprising MCb ^+^ spp.

Figure S2. Phylogenomic analyses of select LB, Bacillaceae, Listeriaceae and Staphylococcaceae spp.

Figure S3. Phylo-mLogo display of site-specific sequence variations in extra-membranous loops between aligned seqs from the MntH C groups MCa, MCb_ie_ and MCb_gut_.

Figure S4. CFnt modeling of *Terribacillus aidingensis* MCb_ie_ A0A285P4W2 responds to mutagenesis like A0A143Y4E1 from *Trichochoccus palustris*.

Figure S5. MCb_gut_ phylogeny in the Enterococcaceae family.

Figure S6. CFpdb modeling shows MCb_gu1_ and MCb_gu2_ structural dynamics differ.

Figure S7. MCb_gu1_ and MCb_gu2_ unrooted phylogeny.

Figure S8. Divergence of archetype Nramp parologs Nramp1 and Dmt1 followed the 1R whole genome that took place in early vertebrates.

Figure S9. The gut microbiota of wild hagfishes may be enriched in MCb ^+^ bacteria.

Table S1. Slc11 carrier nomenclature, taxonomic distribution (representative taxa) and origin.

Table S2. Mutants produced in this study.

## Author Contributions

M.C. is sole author.

## Data Availability Statement

The models and sequences used in this study are included in Appendix A.

## Acknowledgments

The providers and caretakers of online services, servers and repositories that were used for this work are gratefully acknowledged. I thank Stella Cellier-Goetghebeur for comments and suggestions to improve this manuscript.

## Appendix A Contains additional data (Accession numbers of the sequences used; 3D model pdb files).

