## Supplemental Figures & Tables for "Emergence of Slc11 clade MCb_gut_: a parsimonious hypothesis for the dawn of Lactobacillales in the gut of early vertebrates"

A

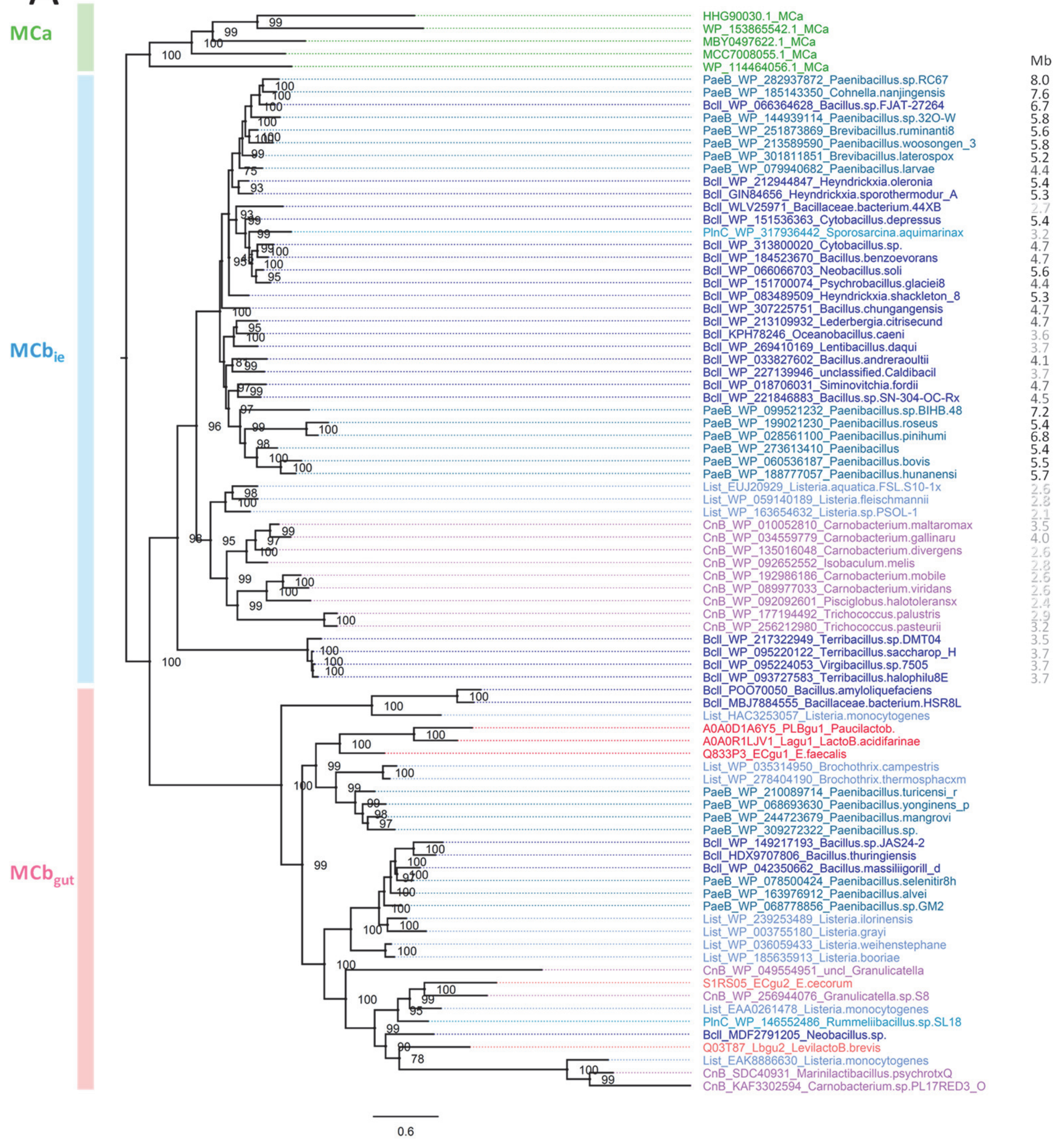


MntH distribution among MCb$_{ie}$$^{+}$ spp. { BACILLALES / LACTOBACILLALES

| | MCb$_{gut}$ | MCb$_{ie}$ | MCa | MA |
|---|---|---|---|---|
| BclL | 8 | 45 | 86 | **140** |
| PlnC | 1 | 8 | 1 | **20** |
| PaeB | 17 | 25 | **205** | 73 |
| List | **14** | 7 | 6 | 0 |
| CrnBc | 7 | **11** | 1 | 0 |
| LctBL | **424** | 0 | 0 | 0 |

Bacillaceae
Planococcaceae
Paenibacillaceae
Listeriaceae
Carnobacteriaceae

B

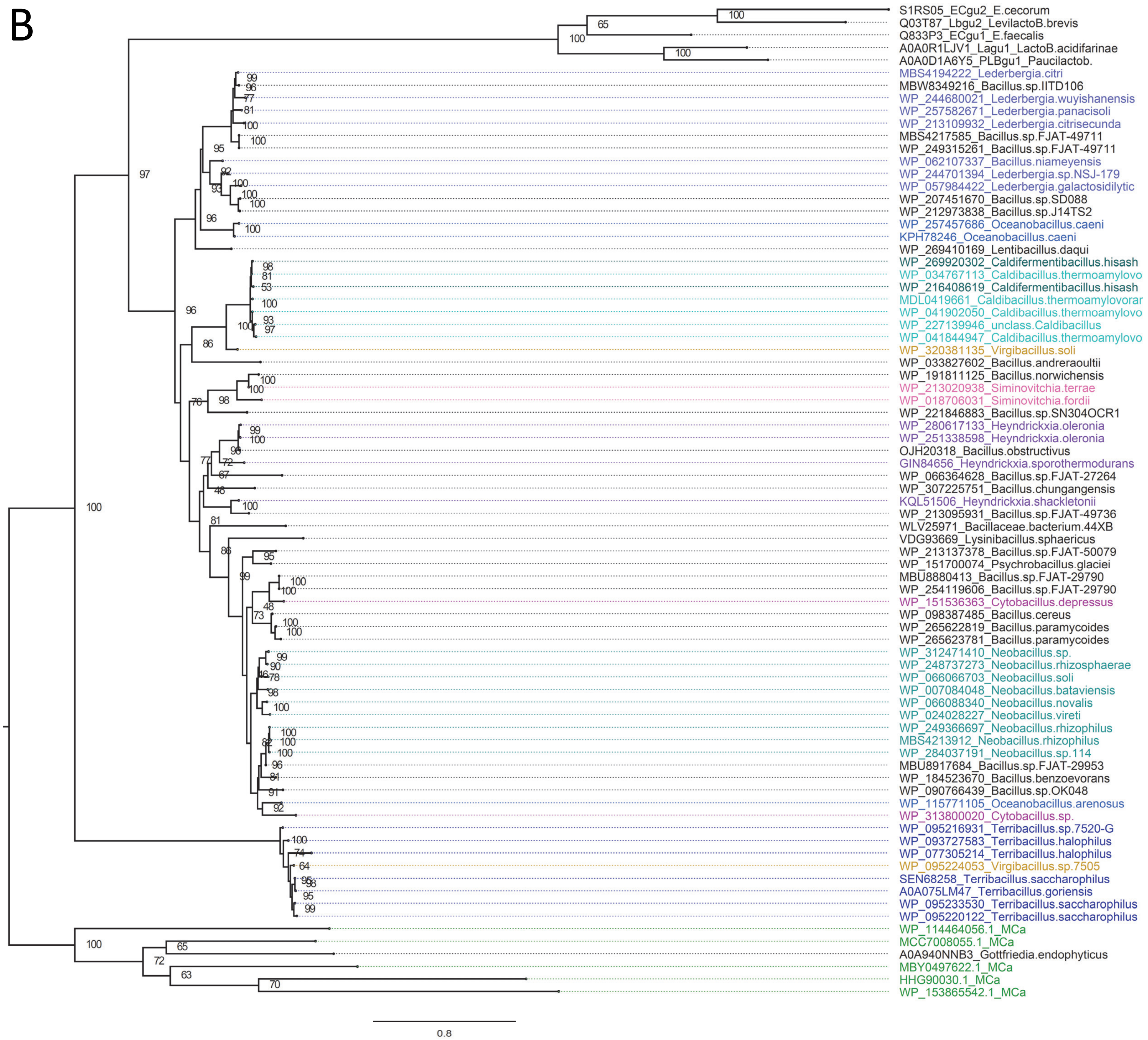
S1RS05_ECgu2_E.cecorum
Q03T87_Lbgu2_LevilactoB.brevis
Q833P3_ECgu1_E.faecalis
A0A0R1LJV1_Lagu1_LactoB.acidifarinae
A0A0D1A6Y5_PLBgu1_Paucilactob.
MBS4194222_Lederbergia.citri
MBW8349216_Bacillus.sp.IITD106
WP_244680021_Lederbergia.wuyishanensis
WP_257582671_Lederbergia.panacisoli
WP_213109932_Lederbergia.citrisecunda
MBS4217585_Bacillus.sp.FJAT-49711
WP_249315261_Bacillus.sp.FJAT-49711
WP_062107337_Bacillus.niameyensis
WP_244701394_Lederbergia.sp.NSJ-179
WP_057984422_Lederbergia.galactosidilytic
WP_207451670_Bacillus.sp.SD088
WP_212973838_Bacillus.sp.J14TS2
WP_257457686_Oceanobacillus.caeni
KPH78246_Oceanobacillus.caeni
WP_269410169_Lentibacillus.daqui
WP_269920302_Caldifermentibacillus.hisash
WP_034767113_Caldibacillus.thermoamylovo
WP_216408619_Caldifermentibacillus.hisash
MDL0419661_Caldibacillus.thermoamylovorar
WP_041902050_Caldibacillus.thermoamylovo
WP_227139946_unclass.Caldibacillus
WP_041844947_Caldibacillus.thermoamylovo
WP_320381135_Virgibacillus.soli
WP_033827602_Bacillus.andreraoultii
WP_191811125_Bacillus.norwichensis
WP_213020938_Siminovitchia.terrae
WP_018706031_Siminovitchia.fordii
WP_221846883_Bacillus.sp.SN304OCR1
WP_280617133_Heyndrickxia.oleronia
WP_251338598_Heyndrickxia.oleronia
OJH20318_Bacillus.obstructivus
GIN84656_Heyndrickxia.sporothermodurans
WP_066364628_Bacillus.sp.FJAT-27264
WP_307225751_Bacillus.chungangensis
KQL51506_Heyndrickxia.shackletonii
WP_213095931_Bacillus.sp.FJAT-49736
WLV25971_Bacillaceae.bacterium.44XB
VDG93669_Lysinibacillus.sphaericus
WP_213137378_Bacillus.sp.FJAT-50079
WP_151700074_Psychrobacillus.glaciei
MBU8880413_Bacillus.sp.FJAT-29790
WP_254119606_Bacillus.sp.FJAT-29790
WP_151536363_Cytobacillus.depressus
WP_098387485_Bacillus.cereus
WP_265622819_Bacillus.paramycoides
WP_265623781_Bacillus.paramycoides
WP_312471410_Neobacillus.sp.
WP_248737273_Neobacillus.rhizosphaerae
WP_066066703_Neobacillus.soli
WP_007084048_Neobacillus.bataviensis
WP_066088340_Neobacillus.novalis
WP_024028227_Neobacillus.vireti
WP_249366697_Neobacillus.rhizophilus
MBS4213912_Neobacillus.rhizophilus
WP_284037191_Neobacillus.sp.114
MBU8917684_Bacillus.sp.FJAT-29953
WP_184523670_Bacillus.benzoevorans
WP_090766439_Bacillus.sp.OK048
WP_115771105_Oceanobacillus.arenosus
WP_313800020_Cytobacillus.sp.
WP_095216931_Terribacillus.sp.7520-G
WP_093727583_Terribacillus.halophilus
WP_077305214_Terribacillus.halophilus
WP_095224053_Virgibacillus.sp.7505
SEN68258_Terribacillus.saccharophilus
A0A075LM47_Terribacillus.goriensis
WP_095233530_Terribacillus.saccharophilus
WP_095220122_Terribacillus.saccharophilus
WP_114464056.1_MCa
MCC7008055.1_MCa
A0A940NNB3_Gottfriedia.endophyticus
MBY0497622.1_MCa
HHG90030.1_MCa
WP_153865542.1_MCa
0.8

**Figure S1.** MCb ($MCb_{ie}$ and $MCb_{gut}$) phylogeny among bacterial families comprising $MCb_{ie}^{+}$ spp. **A.** MntH distribution among $MCb_{ie}^{+}$ spp. (right) was established using PHI-Blast searches and sequence similarity clustering (95% aa id). Details of MCa, $MCb_{ie}$ and $MCb_{gut}$ sequences are provided in Appendix 1. MCb phylogeny (left) was determined using IQ-Tree and 328 PI sites, the substitution model EHO, ML estimate of a.a. state frequency, free rate model of variation among sites with 7 categories. The statistical significance of each node is indicated together with the scale (nb substitution per site). $MCb_{ie}$ and $MCb_{gut}$ sister groups are highlighted, and the size (in megabases) of the genome linked to each $MCb_{ie}$ sequence analyzed is shown next to the spp. names, color-coded as indicated. **B.** Detailed phylogeny of $MCb_{ie}$ among Bacillaceae. Sequence clusters representing established genera are shown using different colors to indicate the corresponding spp. names. IQ-Tree phylogeny was established using 308 PI sites, the substitution model UL3, ML estimate of a.a. state frequency, free rate model of variation among sites with 6 categories and rooted with 5 MCa seqs. The statistical significance of each node is indicated together with the scale (nb substitution per site).

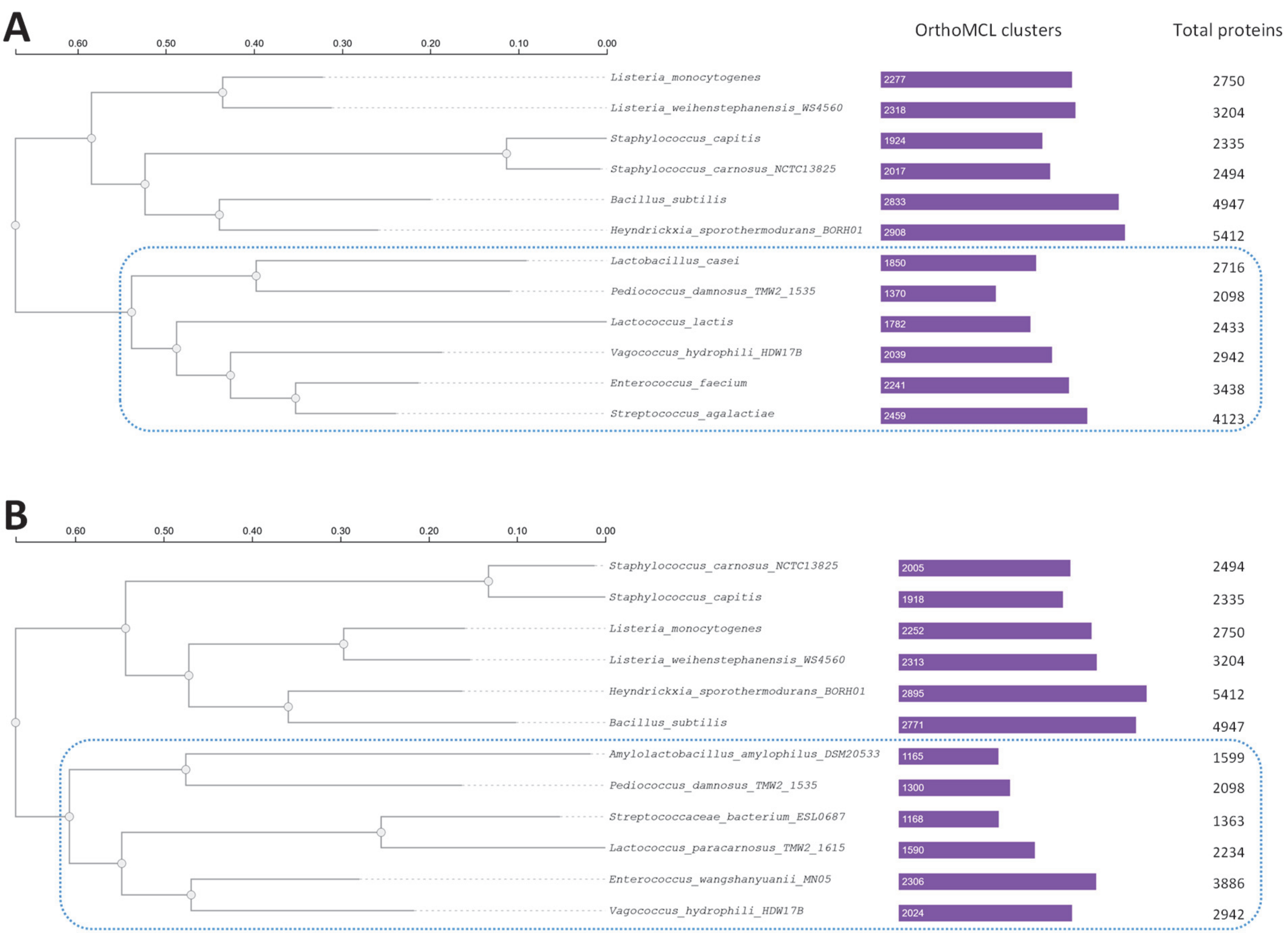


**Figure S2**. Phylogenomic analyses (FastTree, LG-CAT) of select LB, Bacillaceae, Listeriaceae and Staphylococcaceae spp. based on **A**, 165, and **B**, 380 single-copy OrthoMCL clusters (e-value $1^{e-2}$) produced by OrthoVenn3 (Sun et al., 2023, PMID: 37114999). Total numbers of annotated proteins and identified orthologous clusters are listed per genome. While the relative positions of *Listeria* and *Staphylococci* spp. vary with the selection of LB spp., *Heyndrickxia* consistently appears closer than *B. subtilis* to the LB clade (blue dots).

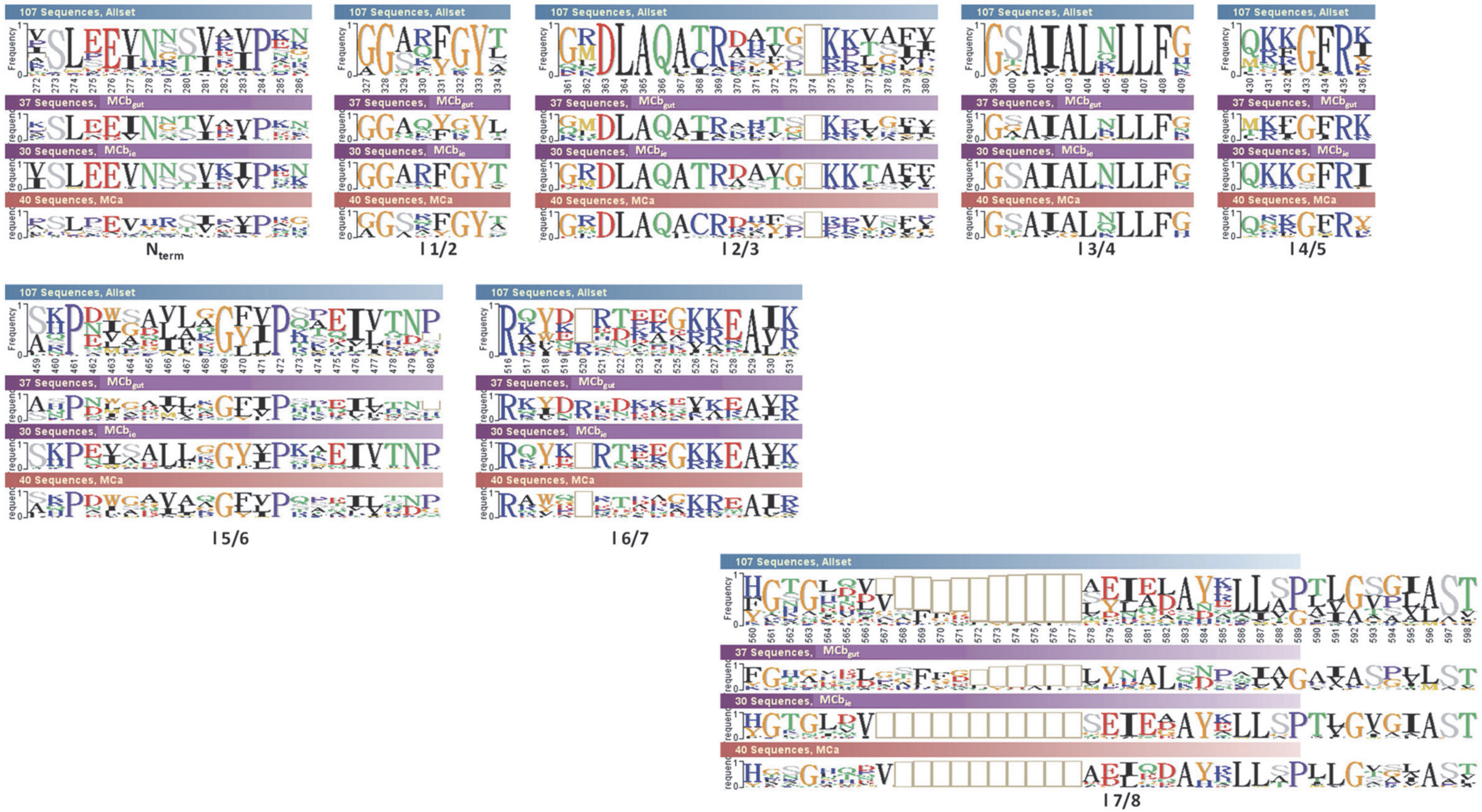

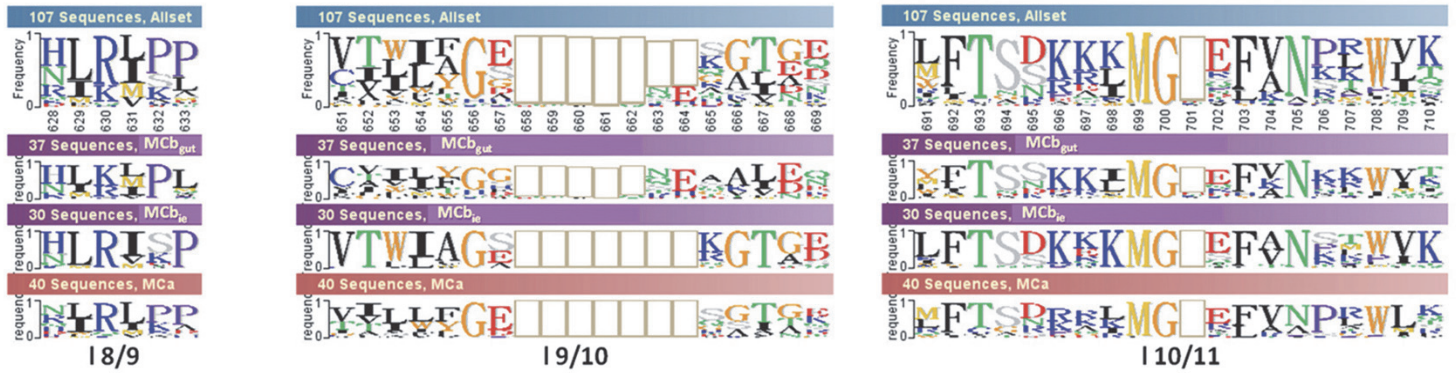


**Figure S3.** Phylo-mLogo display of site-specific sequence variations in extra-membranous loops between aligned seqs from the MntH C groups MCa, $MCb_{ie}$ and $MCb_{gut}$. $MCb_{gut}$-specific sequence divergence is notably prominent in loops (l) 7/8, l 9/10, l 6/7 and l 4/5.

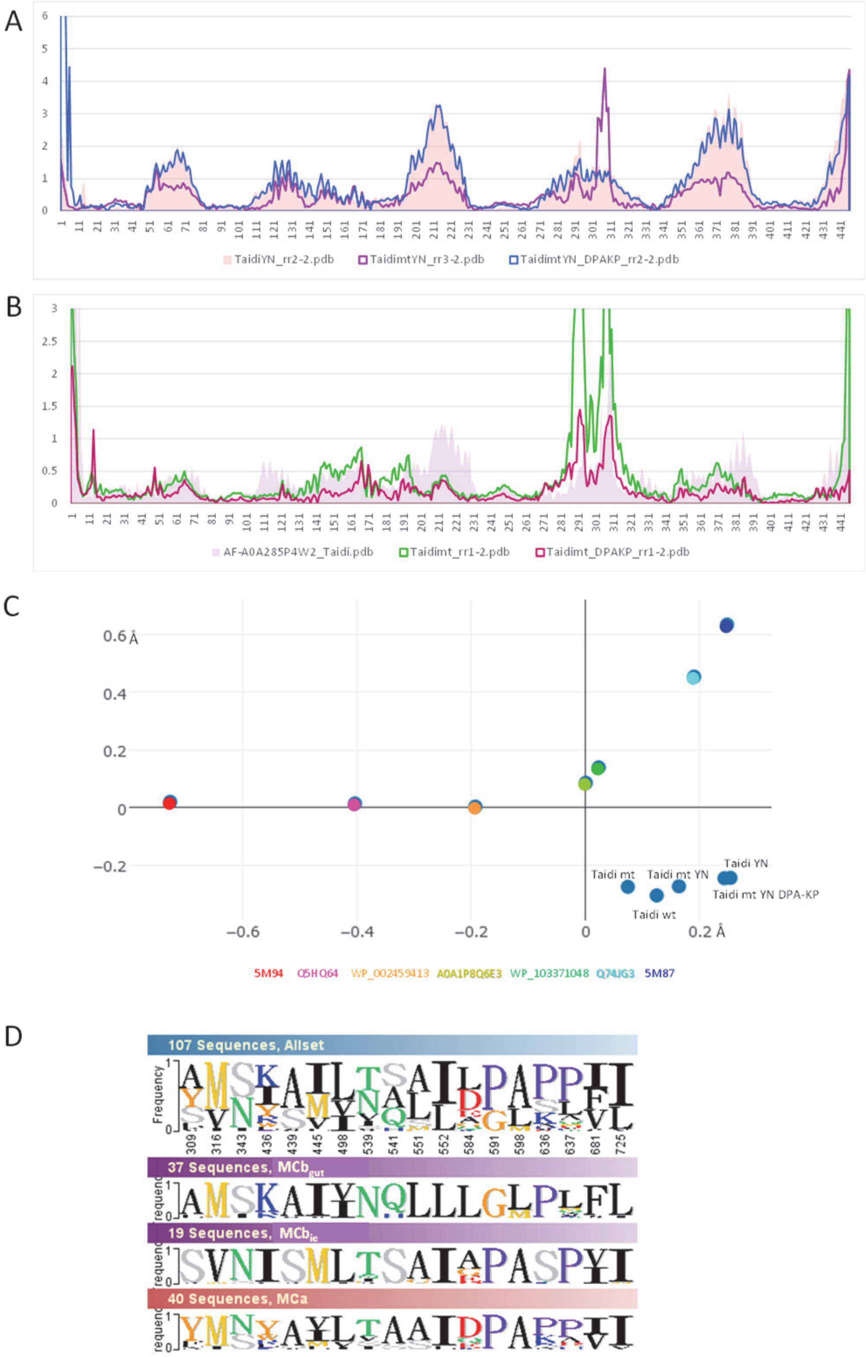


**Figure S4.** CFnt modeling of *Terribacillus aidingensis* $MCb_{ie}$ A0A285P4W2 responds to mutagenesis like A0A143Y4E1 from *Trichococcus palustris*. The 4 panels present pRMSDs relative to native A0A285P4W2 CFnt model (**A,B**), 3D correspondence of native and mutant A0A285P4W2 models with the solved structures 5M87 and 5M94 plus a series of predicted $MCb_{gut}$ conformers (Cellier M., 2023, PMID: 37894758) (**C**) and Phylo-mLogo display of the sites mutated in A0A285P4W2_mt (**D**). **A.** h3 YN mutation-induced conformation shift is frustrated by compound mutagenesis (15 mutations mimicking $MCb_{gut}$ evolutionary rate-shits, D) and restored by reverting of 5 of the 15 sites co-assayed (sites #584, 591, 598, 636, 637, panel D). **B.** Compound mutagenesis induces slight deviation from native A0A285P4W2 model, counter-acted by reverting of 5 of the co-assayed sites. **C.** 3D correspondence of the A0A285P4W2 models studied. **D.** The majority of sites distinguishing $MCb_{gut}$ from $MCb_{ie}$ also demonstrates $MCb_{ie}$ closeness to MCa.

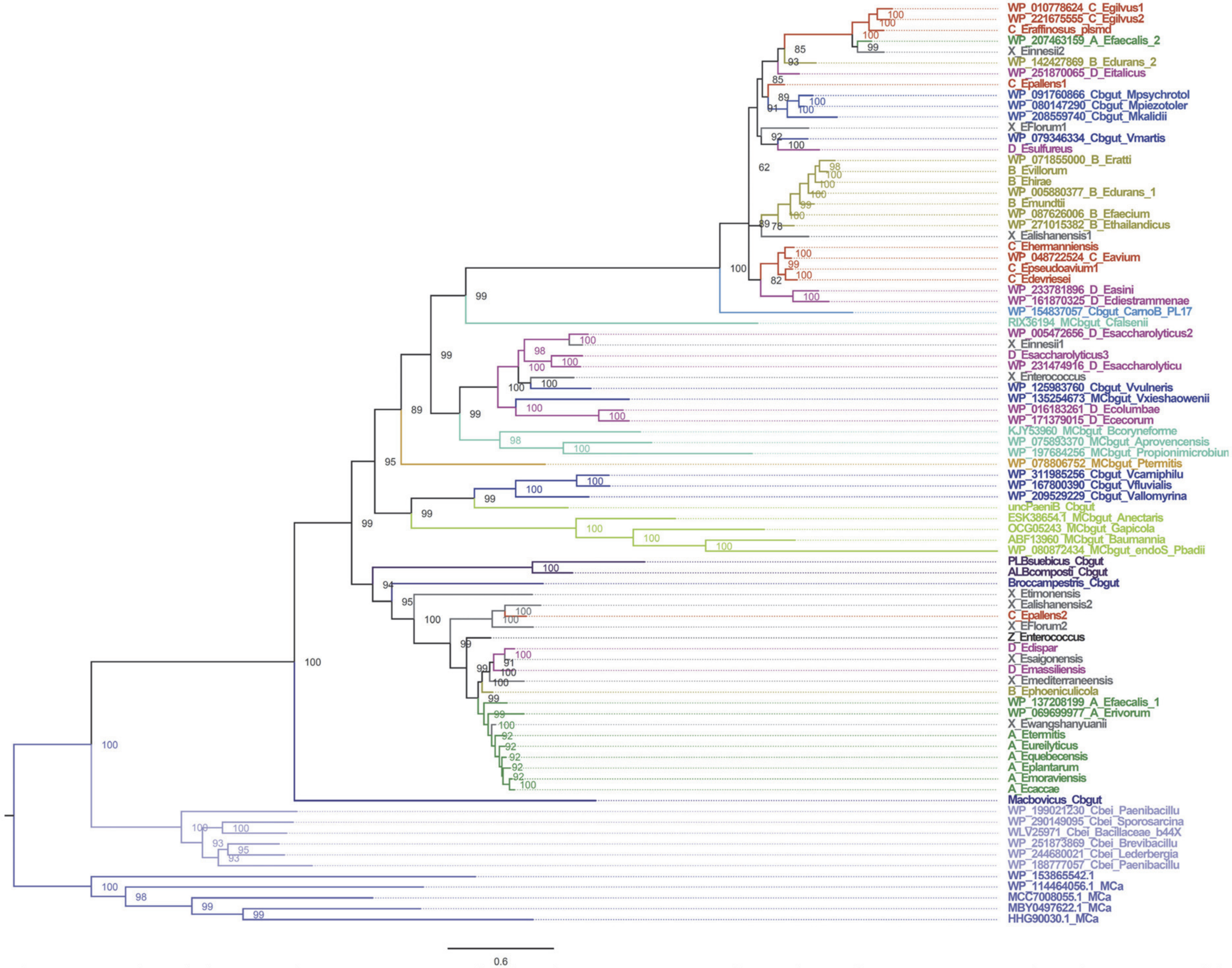


**Figure S5**. $MCb_{gut}$ phylogeny in the Enterococcaceae family. The IQ-Tree presented was obtained using 336 PI sites, the substitution model EX-EHO, ML estimate of a.a. state frequency, free rate model of variation among sites with 12 categories and rooted with 5 MCa seqs. The statistical significance of each node is indicated together with the scale (nb substitution per site). Sequence names are color-coded to distinguish either clades of *Enterococcus* (A, B, C, D) or select genera e.g., *Vagococcus*, *Marinilactibacillus*. Some branches of the tree were colored to highlight sequence clusters of biological significance: for instance, seqs from *Enterococcus* clades A and B form tighter clusters than *Enterococcus* clades C and D, and distinct seq clusters resulting from horizontal transfer to Actinobacteria or Enterobacteria spp. are highlighted.

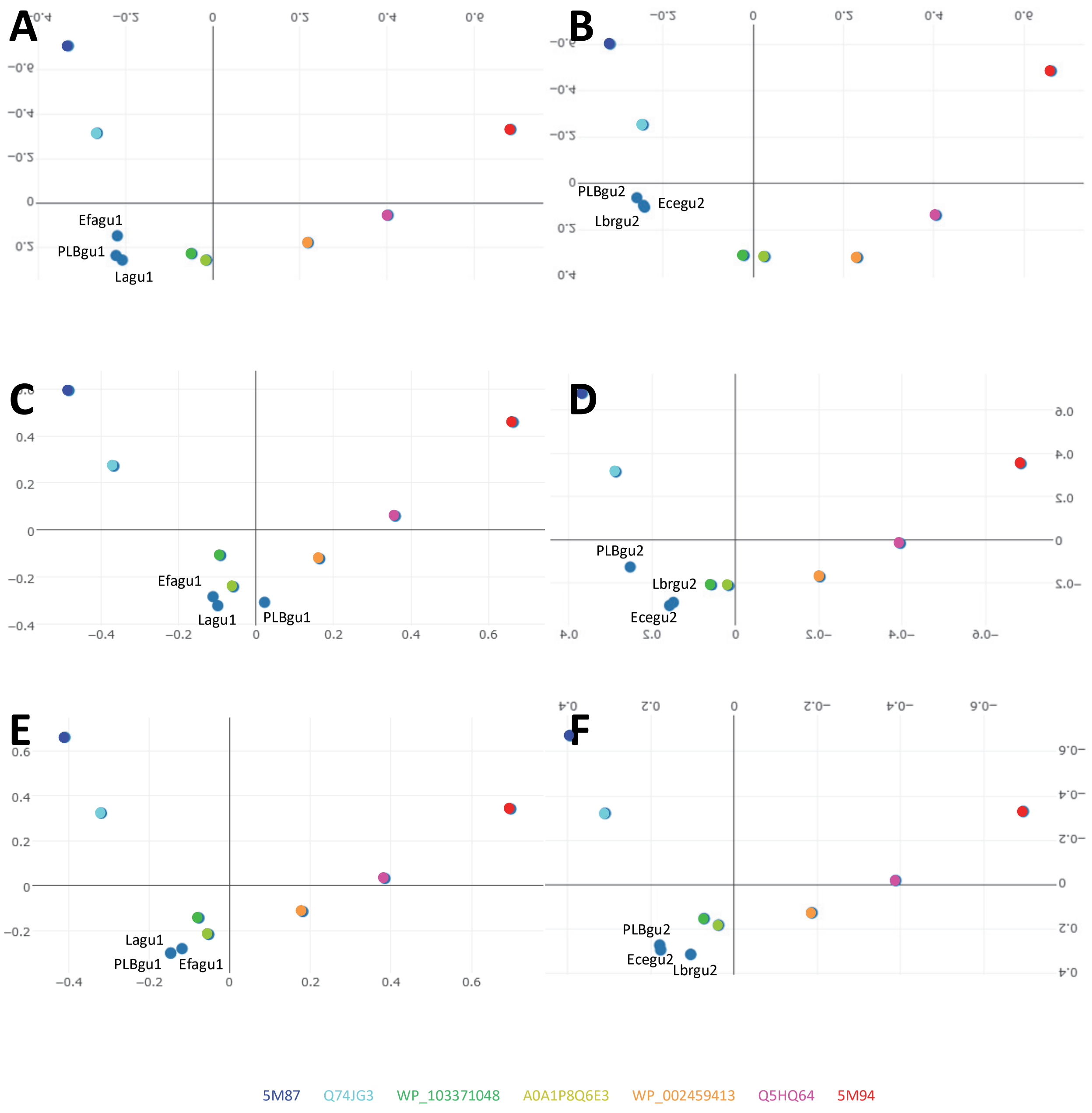


**Figure S6**. CFpdb modeling shows $MCb_{gu1}$ and $MCb_{gu2}$ structural dynamics differ. 3D correspondence analyses (Dali all against all) between a series of $MCb_{gut}$ model conformers (Cellier M., 2023, PMID: 37894758) plus solved structures (5M87, 5M94) and CFpdb models of $MCb_{gu1}$ and $MCb_{gu2}$ templates either native (A, B), mutated to induce carrier forward transition of $PLB_{gu1}$ (C, D), or further reciprocally mutated at sites distinguishing $MCb_{gu1}$ from $MCb_{gu2}$. (E, F).

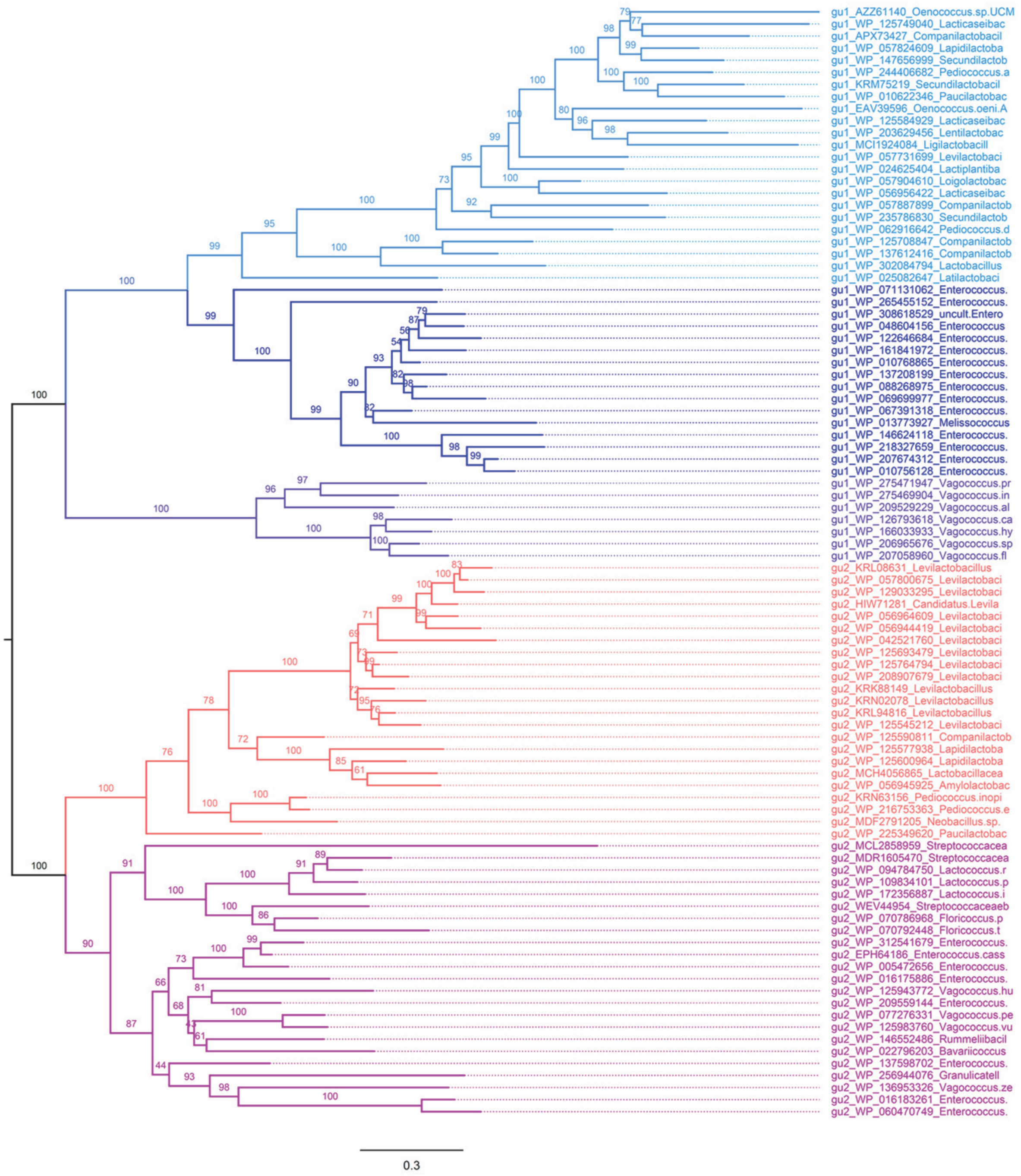


**Figure S7.** $MCb_{gu1}$ and $MCb_{gu2}$ unrooted phylogeny. A set of 285 PI sites was used for IQ-Tree analysis with the substitution model UL3, ML estimate of a.a. state frequency, free rate model of variation among sites with 5 categories. The statistical significance of each node is indicated together with the scale (nb substitution per site).

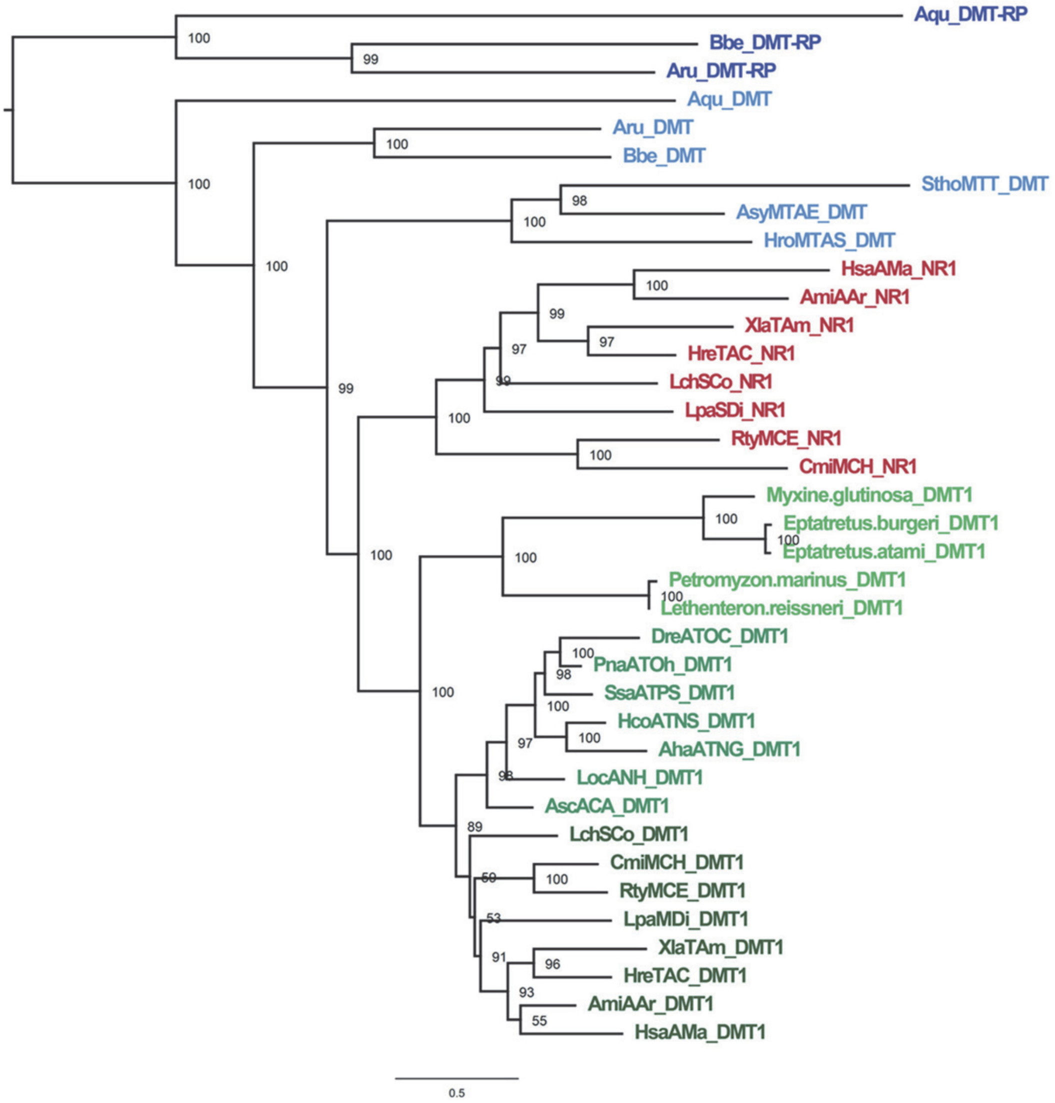


| | Nramp/Dmt-related protein | Nramp/Dmt | Nramp1 | Dmt1 |
|---|---|---|---|---|
| Deuterostoma | | | | |
| Echinodermata | + | + | - | - |
| Urochordata | - | + | - | - |
| Cephalochordata | + | + | - | - |
| Vertebrata | | | | |
| Cyclostomata | - | - | - | + |
| Gnathostomata | | | | |
| Chondrichthyes | - | - | + | + |
| Osteichthyes | | | | |
| Actinopterygii | - | - | - | + |
| Sarcopterygii | - | - | + | + |

**Figure S8.** Divergence of archetype Nramp parologs Nramp1 and Dmt1 followed the 1R whole genome duplication that took place in early vertebrates (Yu et al., 2024, PMID: 38216617, Marletaz et al., 2024, PMID: 38262590). Phylogeny of Nramp/Dmt homologs from basal Deuterostoma [Echinoderms (Aqu), Cephalochordates (Bbe, Aru) and Urochordates (SthoMTT, AsyMTAE, HroMTAS)] and vertebrates [Cyclostomata (Myxine, Eptatretus, Petromyzon, Lethenteron) and Gnathostomata [Chondrichthyes (RtyMCE, CmiMCH) and [Osteichthyes [Actinopterygii (LchSCo, AhaATNG, HcoATNS, DreATOC, PnaATOh, SsaATPS, LocANH, AscACA) and Sarcopterygii (LpaSDi, LchSCo, HreTAC, XlaTAm, AmiAAr, HsaAMa)]]]]. The IQ-tree presented used 313 PI sites sites, the substitution model UL3, ML estimate of a.a. state frequency, free rate model of variation among sites with 7 categories and was rooted using DMT-RP seqs (Sassa et al., 2021, PMID: 33846549). The statistical significance of each node is indicated together with the scale (nb substitution per site). Inset, current estimate of *Nramp/Dmt* complement among Deuterostoma.

SRA TBN hits (word size 3)

*Eptatretus stoutii* GUT CONTENTS

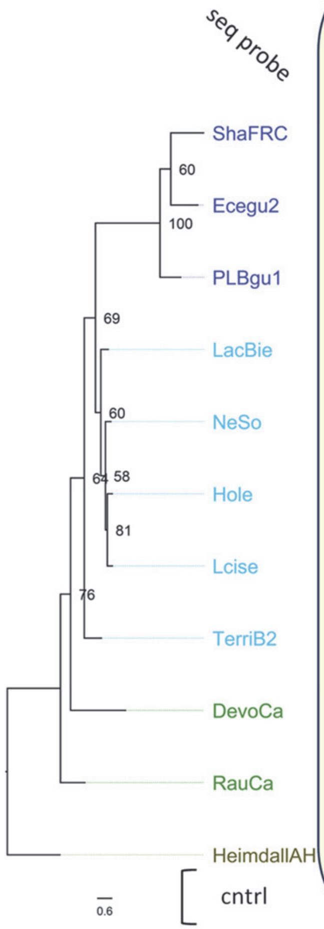


| seq probe | WILD Expect 1e-02 Total | %id range | >50% id | WILD Expect 1e-03 Total | %id range | >50% id | WILD Expect 1e-03 Total | %id range | >50% id | WILD Expect 1e-05 Total | %id range | >50% id | CAPTIVE Expect 1e-05 Total | %id range | >50% id | CAPTIVE Expect 1e-07 Total | %id range | >50% id |
|---|---|---|---|---|---|---|---|---|---|---|---|---|---|---|---|---|---|---|
| ShaFRC | 2 | **68** 48 | 1 | 0 | - | 0 | 14 | 63 45 | **12** | **12** | 63 47 | **11** | 50 | 48 40 | 0 | 22 | 46 41 | 0 |
| Ecegu2 | 0 | - | 0 | 0 | - | 0 | 13 | **69** 48 | 10 | 10 | 69 48 | 7 | 36 | **58** 51 | 36 | 24 | 56 51 | 24 |
| PLBgu1 | 0 | - | 0 | 0 | - | 0 | **15** | **71** 46 | 11 | 10 | **71** 52 | 10 | 33 | 48 42 | 0 | 1 | 47 | 0 |
| LacBie | **4** | **64** 47 | **3** | **1** | **64** | **1** | **16** | 66 48 | **13** | 7 | 66 52 | 7 | **94** | **57** 46 | **76** | **35** | **57** 48 | 33 |
| NeSo | 1 | 54 | 1 | 0 | - | 0 | 12 | 60 45 | 9 | 6 | 60 50 | 6 | 70 | 55 42 | 28 | 28 | 54 49 | 20 |
| Hole | 1 | 54 | 1 | 0 | - | 0 | 13 | 64 47 | 8 | 6 | 64 54 | 6 | **91** | 56 44 | **75** | **35** | 56 51 | **35** |
| Lcise | 1 | 54 | 1 | 0 | - | 0 | 11 | 60 45 | 6 | 6 | 60 48 | 5 | 59 | 52 40 | 16 | 28 | 52 44 | 7 |
| TerriB2 | **4** | 47 42 | 0 | 1 | 46 | 0 | 14 | 64 46 | **12** | 9 | 64 51 | 9 | 50 | 52 44 | 31 | 28 | 52 47 | 19 |
| DevoCa | 1 | 45 | 0 | 0 | - | 0 | 8 | 55 43 | 3 | 6 | 53 47 | 2 | 32 | 48 45 | 0 | 9 | 48 | 0 |
| RauCa | 3 | 54 37 | 1 | 0 | - | 0 | 9 | 64 42 | 3 | 5 | 64 45 | 3 | 63 | 55 45 | 38 | **35** | 55 49 | 28 |
| HeimdallAH | SRA: ERS5470882 | | | | | | SRA: ERS5470883 | | | | | | SRA: ERS5471345 | | | | | |
| cntrl | | | | **29** | **100** 51 | **29** | | | | **32** | **100** 40 | **31** | | | | **355** | **85** 50 | **355** |

**Figure S9**. The gut microbiota of wild hagfishes may be enriched in $MCb_{gut}^{+}$ bacteria. SRAs linked to the marine fish gut microbiome sequencing project (Bioproject PRJNA720542) were screened by TBlastN analysis using phylogenetically defined probes representative of MCa, $MCb_{ie}$ and $MCb_{gut}$ clades. For each probe the total number of hits, their range of sequence identity and the number of hits showing more than 50% identity are indicated. The top 2 values in each column are bolded. All probes were assayed at two stringencies by varying the value of expected threshold (number of chance matches). Probes: Sha FRC (WP_011276123.1), Ecegu2 (S1RS05), PLBgu1 (A0A0D1A6Y5), LacBie (EUJ20929), NeSo (WP_066066703), Hole (WP_212944847), Lcise (WP_213109932), TerriB2 (WP_095216931), DevoCa (HHG90030), RauCa (WP_114464056); out group, HeimdallAH (MAG Candidatus *Heimdallarchaeota* archaeon, DAHXPE010000008.1). Control probes: *Eptatretus atami* Nramp2/Dmt1 (JAXMNL010000005, SRA: ERS5470882, SRA: ERS5470883), *Escherichia coli* MntH A (WP_000186386, SRA: ERS5471345). Note, for SRA: ERS5470883, only one hit obtained with *E. atami* probe (40% id) also cross-reacted with *E. faecalis* $MCb_{gu1}$ (WP_0137208199, 50% id) and with *E. coli* MA (98% id); top *E. faecalis* $MCb_{gu1}$ hits (>55% id) did not cross react with *E. coli* MA. In contrast, all hits obtained from SRA: ERS5470882 with MCb probes ($MCb_{ie}$ and $MCb_{gut}$) showed higher conservation with *E. atami* probe while *E. coli* MA probe returned no hits.

Table S1. Slc11 carrier nomenclature, taxonomic distribution (representative taxa) and origin[1]

| Name | Symbol | PROKARYOTE | pre-EUKARYOTE | EUKARYOTE |
|---|---|---|---|---|
| MntH[2] B [***B***efore aerobiosis] | MB | Bacteroidetes, Firmicutes, DeltaPB[4] | | |
| MntH A [***A***erobes] | MA | GammaPB, Firmicutes Actinobacteria | | |
| MntH H [***H***yperthermophiles] | MH | | TACK[5]/Asgard Archaea Deinococcus/Thermus GammaPB | |
| Archetype Nramp[3] | aN | | | |
| | aN-I | | | SAR[6], ARCHAEPLASTIDA |
| | aN-II | | | FUNGI, AMOEBA, RHODOPHYTA METAZOA |
| Prototype Nramp | pN | | | |
| | pN-I | | | SAR, FUNGI, ARCHAEPLASTIDA |
| | pN-II | | | GLAUCOPHYTA, RHODOPHYTA AMOEBA |
| MntH C [Eukaryotic ***C***ell-derived] | MC | | | |
| MntH Ca [pN-I-derived] | MCa | | | Firmicutes, AlphaPB Bacteroidetes, Cyanobacteria |
| MntH Cb [MCa-derived] | MCb | | | Firmicutes |
| MntH Cg [MCa-derived] | MCg | | | GammaPB, BetaPB |
| MntH CaU [pN-II-derived] | MCaU | | | Bacteroidetes |

[1]Cellier M., 2012, PMID: 23046654
[2]prokaryotic proton ($H^+$)-dependent manganese ($Mn^{2+}$) transporter
[3]eukaryotic natural resistance-associated macrophage protein, a.k.a. divalent metal transporter (Dmt)
[4]Proteobacteria
[5]Thaumarchaeota (now Nitrososphaerota), Aigarchaeota, Crenarchaeota (now Thermoproteota), and Korarchaeota
[6]supergroup including Stramenopiles, Alveolates, Rhizarians

TABLE S2 Mutants produced in this study

| | |
|---|---|
| **Figure 5:** | |
| Y4E1mt | S57A, V64M, W183K, S186A, M192I, L237V, F277N, S279Q, I290L, D310L, P317G, A324L, K360P, P361L, I398F, I440L |
| T065mt | V58M, N85S, I177K, S180A, M186I, L231V, F271N, S273Q, M283L, I284L, D304L, P311G, A318L, E354P, P355L, I392F, I434L |
| **Figure 6:** | |
| JG317 | L42S, M49V, S76N, K168I, A171S, I177M, V228L, N269T, Q271S, L281A, L282I, M312A, I319P, L327A, P363S, L364P, F408V, L450I |
| Q6E317 | L38S, M45V, S72N, K164I, A167S, I173M, I218L, N259T, Q261S, L271A, L272I, L297A, G304P, L311A, P347S, L348P, F387V, L429I |
| Q17pdb | L46S, M53V, S80N, K172I, A175S, I181M, V226L, N267T, Q269S, L279A, L280I, L306A, G313P, M320A, P356S, N357P, F396V, L438I |
| V1J217 | L42S, M49V, S76N, K168I, A171S, I177M, V222L, N263T, Q265S, L275A, L276I, L302A, G309P, M316A, P352S, N353P, F392V, L434I |
| WM917 | L39S, M46V, S73N, K165I, A168S, I174M, I219L, N260T, Q262S, L272A, L273I, L299A, G306P, M313A, P349S, N350P, F389V, L431I |
| **Figure 7:** | |
| Y4E1nt_VNT | A142Y, G146N, A239T, M242V, H244Y |
| Y4E1nt_VY | M242V, H244Y |
| Y4E1nt_YN | A142Y, G146N |
| Y4E1ntrVN | A142Y, G146N, M242V, H244Y |
| **Figure 8:** | |
| Y4E1ntYN_396A | A142Y, G146N, Q396A |
| Y4E1ntYN_396G | A142Y, G146N, Q396G |
| Y4E1ntYN_396N | A142Y, G146N, Q396N |
| Y4E1ntYN_396S | A142Y, G146N, Q396S |
| Y4E1ntYN66A | P66A, A142Y, G146N |
| Y4E1ntYN68G | N68G, A142Y, G146N, |
| Y4E1ntYN66A68G | P66A, N68G, A142Y, G146N, |
| TpalY4E1mtIL | Y4E1mt M137I, I141L |
| TpalY4E1mtILYN | TpalY4E1mtIL A142Y, G146N |
| TpalY4E1mtYN | Y4E1mt A142Y, G146N |
| Y4E1ntILYN | Y4E1nt M137I, I141L, A142Y, G146N, |
| Tpal_Y4E1mtAYN | TpalY4E1mtYN L324A |
| Tpal_Y4E1mtDPYN | TpalY4E1mtYN L310D, G317P |
| Tpal_Y4E1mtDPAYN | TpalY4E1mtYN L310D, G317P, L324A |
| **Figure 9:** | |
| Y4E1mtDPAYN_I | Tpal_Y4E1mtDPAYN F398I |
| Y4E1mtDPAYN_KP | Tpal_Y4E1mtDPAYN P360K, L361P |
| Y4E1mtDPAYN_KPI | Tpal_Y4E1mtDPAYN P360K, L361P, F398I |
| Y4E1mtYN_KP | TpalY4E1mtYN P360K, L361P |
| Tpal_Y4E1mtDPA | Y4E1mt L310D, G317P, L324A |
| Y4E1mt_DPAKP | Y4E1mt L310D, G317P, L324A, P360K, L361P |
| Y4E1mt_DPAKPI | Y4E1mt L310D, G317P, L324A, P360K, L361P, F398I |
| Y4E1mt_KP | Y4E1mt P360K, L361P |
| Y4E1mtDPAI | Y4E1mt L310D, G317P, L324A, F398I |
| **Figure S4:** | |
| Taidimt | V45A, I52M, N79S, W171K, M180I, L225V, T265N, S267Q, A277L, I278L, E298L, P305G, A312L, K348P, P349L, I386F, I428L |
| TaidiYN | A130Y, G134N |
| TaidimtYN | Taidimt A130Y, G134N |
| TaidimtYN_DPAKP | TaidimtYN L298D, G305P, L312A, P348K, L349P |
| Taidimt_DPAKP | Taidimt L298D, G305P, L312A, P348K, L349P |
| **Figure 12:** | |
| PLBgu1_TY | M233T, H235Y |
| PLBgu1_TYG | PLBgu1_TY D56G |
| PLBgu1_TYG7T | PLBgu1_TYG N279T |
| 1_TYG7TtIFI | PLBgu1_TYG7T V52I, V179F, S335I |
| 1_TYG7TtIFILA | 1_TYG7TtIFI Q334L, T337A |
| 1_TYG7TtIFILA_AN | 1_TYG7TtIFILA D131A, R366N |
| 2a_ANt3AGIi2G | 1_TYG7TtIFILA_AN L119G, I127A, I135G, S138I |
| Lagu1_TYG7T_IFI8LA_AN3AGIi2G | V56I, D60G, L123G, I131A, D135A, I139G, S142I, V183F, M237T, H239Y, N283T, Q338L, S339I, T341A, R369N |
| Efagu1_TYG7T_IFI8LA_AN3AGIi2G | V49I, D53G, L116G, I124A, D128A, V132G, G135I, V176F, M230T, H232Y, N276T, Q331L, N332I, T334A, R362N |
| 2a_ANt3AGIi2G_6iirs | 2a_ANt3AGIi2G I156L, F222T, V313A, S343T, I387L, E388D |
| Lagu1_TYG7T_IFI8LA_AN3AGIi2G_6iirs | Lagu1_TYG7T_IFI8LA_AN3AGIi2G I160L, Y226T, V317A, A347T, I391L, E392D |
| Efagu1_TYG7T_IFI8LA_AN3AGIi2G_6iirs | Efagu1_TYG7T_IFI8LA_AN3AGIi2G I153L, F219T, V310A, S340T, V384L, E385D |
| 2_TYG7TIFI8LAtAN3GAGI | V54I, D58G, L121G, I129A, D133A, V137G, G140I, V181F, M234T, H236Y, N280T, Q335L, N336I, T338A, R367N |
| Lbrgu2_TYG7TIFI8LAtAN3GAGI | V51I, D55G, L118G, I126A, D130A, V134G, G136I, V178F, M231T, H233Y, N277T, Q333L, N334I, T336A, R365N |
| Ecegu2_TYG7TIFI8LA_AN3GAGI | V50I, D54G, L117G, V125A, D129A, I133G, S136I, V177F, M230T, H232Y, N276T, Q331L, N332I, T334A, R363N |
| 2_TYG7TIFI8LAtAN3GAGI_6ii | 2_TYG7TIFI8LAtAN3GAGI L158I, T223F, A314V, T344S, L388I, D389E |
| Lbrgu2_TYG7TIFI8LA(t)AN3GAGI_6iirs | Lbrgu2_TYG7TIFI8LAtAN3GAGI L155I, T220F, A312V, T342S, L386I, D387E |
| Ecegu2_TYG7TIFI8LA_AN3GAGI_6iirs | Ecegu2_TYG7TIFI8LA_AN3GAGI L154I, T219F, A310V, A340S, L384I, D385E |
